# Targeting the deNEDDylating enzyme NEDP1 to ameliorate ALS phenotypes through Stress Granules dissolution

**DOI:** 10.1101/2023.01.06.522988

**Authors:** Toufic Kassouf, Rohit Shrivastava, Igor Meszka, Aymeric Bailly, Jolanta Polanowska, Helene Trauchessec, Jessica Mandrioli, Serena Carra, Dimitris P. Xirodimas

**Author notes:** Equal contribution (Toufic Kassouf, Rohit Shrivastava).

## Abstract

In Amyotrophic Lateral Sclerosis (ALS) motor neuron disease, mutations in proteins that upon stress localize within cytoplasmic protein inclusions called Stress Granules (SGs), are linked to the formation of aberrant inclusions, which are related to neuronal cell death. By combining studies in human cells and *C. elegans* including the use of Nanobodies, we found that inhibition of NEDP1, the enzyme responsible for the processing and deconjugation of the Ubiquitin-like molecule NEDD8 from substrates, promotes the elimination both of physiological and pathological SGs. The hyper-NEDDylation of Poly-(ADP-ribose) polymerase-1 enzyme upon NEDP1 inhibition compromises PAR production and is a key mechanism for the observed SG phenotype. Importantly, the above-described effects are related to improved cell survival in human cells, and in *C. elegans*, NEDP1 deletion ameliorates ALS-phenotypes related to animal motility. Our studies reveal NEDP1 as potential therapeutic target for ALS, based on the elimination of aberrant SGs.

## Introduction

Organisms are constantly exposed to environmental stresses that cause protein damage and the generation of RNA-protein inclusions (proteotoxic stress). A series of sophisticated mechanisms that constitute the so-called Protein Quality Control (PQC) system ensure the detection, repair and/or elimination of damaged proteins and dissolution of inclusions in order to maintain protein homeostasis (proteostasis) (Amm et al., 2014; Chen et al., 2011). Defects in the PQC system lead to the accumulation of aberrant inclusions, which are linked to human diseases, including neurodegeneration and cancer and are also regarded as the hallmark of aging (Hipp et al., 2014, 2019; Pilla et al., 2017).

Recent evidence suggests that two types of proteinaceous inclusions can be induced upon proteotoxic stress conditions: misfolded aggregates and biomolecular condensates that often contain proteins and nucleic acids (Yoo et al., 2022). Although both depend on the PQC system for dispersal, proteinaceous aggregates and condensates show different properties. Aggregates are static structures that require disaggregation or digestion mechanisms for their removal. Instead, stress-induced condensates are more dynamic and form through principles of liquid-liquid phase separation (LLPS) (Alberti and Hyman, 2021; Yoo et al., 2022). One typical example is represented by Stress Granules (SGs), which are rapidly formed under conditions of heat shock or oxidative stress, as part of the response to reduce protein synthesis during proteotoxic stress and to protect mRNA from degradation. Their formation is initiated by the assembly of core proteins around stalled mRNAs, followed by their nucleation mainly with RNA binding proteins that form the shell. Upon stress alleviation, SGs are disassembled, allowing protein synthesis resumption and cell recovery (Mandrioli et al., 2020; Wheeler et al., 2016).

However, defects at the level of SG disassembly can occur and condensates can mature into a less dynamic state that resembles proteinaceous aggregates. The aberrant conversion of SGs from liquid-like dynamic condensates into aggregated-like structures has been associated to the progression of several neurodegenerative diseases, such as the motor neuron disease Amyotrophic Lateral Sclerosis (ALS) or frontotemporal dementia (FTD) (Advani and Ivanov, 2020). Different mechanisms have been proposed for such defects in SG disassembly. First, mutations in genes that encode for SG proteins (e.g. TIA1, FUS, TDP-43) usually occur in the Intrinsically Disordered Region (IDRs), which is a key regulatory module of phase separation and SG dynamics. These mutations confer higher aggregation propensities to these proteins and promote the maturation of liquid-like SGs into aberrant SGs that can further evolve with time into proteinaceous inclusions that persist in the cytoplasm. Second, failure of the PQC system to clear mutated and misfolded SG proteins, or defective ribosomal products (DRiPs), can cause their accumulation within SGs, promoting their conversion into a solid-like aberrant state (Ganassi et al., 2016; Turakhiya et al., 2018). As a result, SGs fail to disassemble upon stress relief, leading to protein synthesis deficiency and contributing to neuronal cell death (Zhang et al., 2019). ALS remains incurable with an average life expectancy of 2-5 years post diagnosis. Acceleration of SG disassembly and elimination of toxic protein inclusions are thus regarded as an attractive approach to potentially re-establish normal neuronal function.

One of the main effectors and regulators of the PQC system is the family of Ubiquitin and Ubiquitin-like molecules (Ubls) such as SUMO and NEDD8. Their covalent modifications on substrate proteins occurs via a three-step process involving E1, E2 and E3 enzymes and are regarded as key regulators of protein oligomerisation and assembly into macromolecular complexes (Abidi and Xirodimas, 2015; Amm et al., 2014; Liebelt and Vertegaal, 2016; Williamson et al., 2013). Several lines of evidence have indicated that Ubiquitination and SUMOylation control the proteome composition and the dynamics of SG assembly and disassembly, even if their role may be dependent on cell-type and applied stress (Bennett and La Spada, 2021; Gwon et al., 2021; Jongjitwimol et al., 2016; Keiten-Schmitz et al., 2020; Marmor-Kollet et al., 2020; Maxwell et al., 2021; Tolay and Buchberger, 2021). These findings further underscore the existence of a tight interplay between the PQC system and SG dynamics. In a siRNA screen for factors that control SG dynamics, components of the NEDD8 pathway were identified (Jayabalan et al., 2016). The role of NEDD8 remains somehow elusive (Meszka et al., 2022), as later studies using short-term chemical inhibition of protein NEDDylation, suggested that the NEDD8 pathway may not be implicated in the control of SG dynamics (Markmiller et al., 2019).

Protein modification with Ubiquitin/Ubls is a reversible process and the extent of substrate modification is finely balanced by the action of deconjugating enzymes (Lange et al., 2022; Williamson et al., 2013). For the NEDD8 pathway there are 2 highly specific deconjugating enzymes, the COP9 signalosome and NEDP1 (Abidi and Xirodimas, 2015; Enchev et al., 2014). NEDP1 also processes NEDD8 into the mature form, before activation by the NEDD8 E1 activating enzyme (NAE) (Meszka et al., 2022). The function of COP9 is established as regulator of the activity of the Cullin-Ring-Ligases (CRLs) through deNEDDylation of Cullins; by contrast, our knowledge on processes controlled by NEDP1 is limited (Meszka et al., 2022; Santonico, 2020). Based on the highly specific activity of NEDP1 to deconjugate NEDD8, understanding the biological role of this enzyme will also assist the elucidation of processes controlled by protein NEDDylation. Indeed, recent studies on NEDP1 identified roles for NEDD8 through modification of non-cullin substrates and the formation of poly-NEDD8 chains as cytoplasmic PQC signal in the DNA damage response (Bailly et al., 2019; Keuss et al., 2019; Meszka et al., 2022; Vijayasimha and Dolan, 2021).

Here, we identify NEDP1 as critical regulator of SGs dynamics. NEDP1 knockout, as well as inhibition of NEDP1 using expression of anti-NEDP1 Nanobody, promotes the disassembly of SGs. Proteome-wide analysis of NEDDylation sites at endogenous level, identified the poly(ADP)-ribosylation polymerase 1 (PARP-1) as NEDP1 substrate. PARP-1 hyper-NEDDylation compromises the generation of PAR conjugates, which are important regulators of SG dynamics (Duan et al., 2019; Jin et al., 2021; McGurk et al., 2018). Critically, we found that NEDP1 inhibition in several *in vitro* systems, including ALS patient-derived fibroblasts, also promotes the elimination of pathogenic SGs. Using *C. elegans* as model organism for ALS, we found that NEDP1 deletion ameliorates ALS phenotypes related to animal motility. Our studies identify NEDP1 as critical regulator of SG dynamics and a potential therapeutic target for ALS.

## Results

### Identification of NEDP1 as regulator of SG dynamics

While Ubiquitin and SUMO conjugation emerge as important regulators of SGs dynamics, the role of NEDD8 in this process has been controversial (Jayabalan et al., 2016; Markmiller et al., 2019). The deNEDDylating enzyme NEDP1 is a critical and highly specific component of the NEDD8 cycle that deconjugates NEDD8 mainly from non-cullin substrates (Meszka et al., 2022). Inhibition of NEDP1 results in the dramatic accumulation of NEDD8 conjugates that amongst other functions control the PQC system in the cytoplasm upon DNA damage. The study of NEDP1 provides a highly specific approach to decipher the role of protein NEDDylation in SGs dynamics.

We generated cell lines stably expressing GFP-G3BP1, the nucleator of SGs, in parental or NEDP1 knockout H6 U2OS cells (Tourrière et al., 2003). Exposure of cells to the SG inducer sodium arsenite (As) caused an accumulation of NEDD8 conjugates, consistent with the idea that the NEDD8 pathway is part of the As stress response (Figure 1A). Importantly, the observed increase in NEDD8 modification strictly depends on the canonical NEDD8 E1 activating enzyme and not on the Ubiquitin system, which can activate and conjugate NEDD8 on substrate proteins under conditions of proteotoxic stress (Hjerpe et al., 2012; Leidecker et al., 2012) (S1A). NEDP1 knockout causes a dramatic increase in protein NEDDylation and no further increase was observed upon As treatment (Figure 1A). Using GFP-G3BP1 as SG marker, we found that NEDP1 deletion accelerated the formation of SGs induced upon As compared to control cells (Figure 1B, C). Similar results were obtained using heat shock as another inducer of SG formation (S1B). The effect is not due to changes in the levels of SGs proteins upon NEDP1 deletion (S2A). In addition, we also assessed the effect of transient inhibition of NEDP1 on SG assembly upon stress. For this, we transiently expressed an anti-NEDP1 Nanobody (Nb9) that displays nM binding affinity and specifically inhibits NEDP1 activity (Abidi et al., 2020). Characterisation *in vitro* and in tissue culture cells, shows that Nb9 potently inhibits both the processing and deconjugating activity of NEDP1, resulting in the accumulation of NEDD8 conjugates to a level similar to that achieved upon NEDP1 knockout (Abidi et al., 2020). Consistent with this notion, expression of Nb9 increases protein NEDDylation and accelerates SG assembly, similarly to what is observed upon deletion of the NEDP1 gene (Figure 1D-F). To better define the role of NEDP1 in SGs assembly, we monitored the size and number of SGs formed upon As using the GFP-G3BP1 cells and live imaging. We found that NEDP1 deletion reduces the size and increases the number of formed SGs upon As (Figure 1G-I, video S1A, B). Similar observations were made when we used endogenous TIA1 as SG marker in control and NEDP1 knockout cells exposed to As (S2B-D). The observed defects in size and number of SGs upon NEDP1 knockout, were rescued upon inhibition of the NEDD8 pathway using specific inhibitors of NAE (MLN4924/NAEi) (Figure 1J, K); this result shows that the observed effects of NEDP1 on SGs are due to the increase in protein NEDDylation.

**Figure 1.**
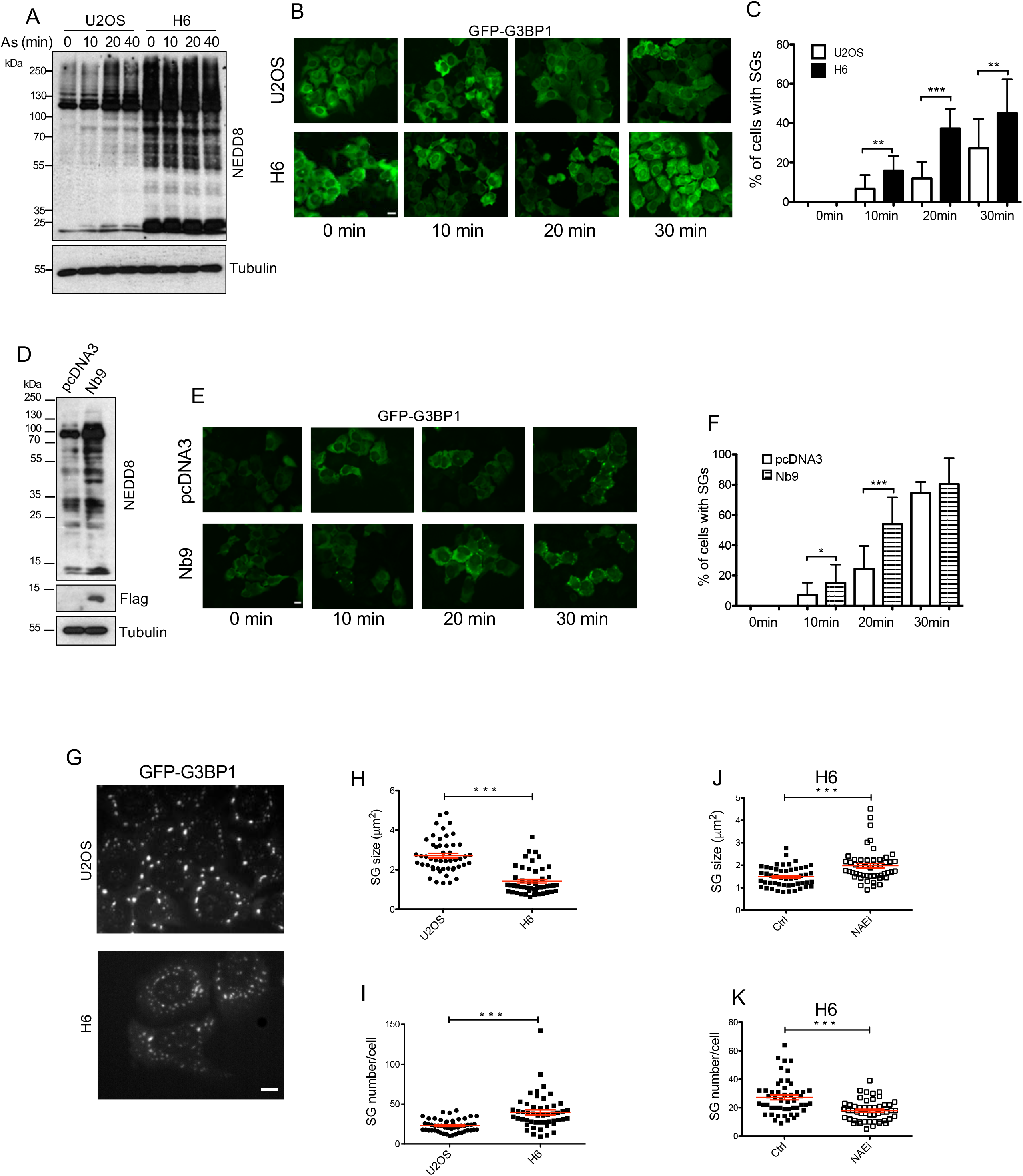
A. Parental and H6 NEDP1 KO U2OS cells were treated with sodium arsenite (As) (0.5mM) then lysed at indicated time points. Protein lysates were analyzed by western blotting using anti-NEDD8 and β-Tubulin antibodies. B. Control and H6 NEDP1 KO U2OS cells stably expressing GFP-G3BP1 were treated with As (0.2mM) then fixed with paraformaldehyde at indicated time points. Scale bars, 10μm. C. Quantification of the experiment performed in B showing the percentage of cells with SGs. Values represents the mean of 3 independent experiments ± SD. D. U2OS cells were transfected with empty (pcDNA3) or Nb9-Flag expressing constructs (2*μ*g). Extracts were analysed by western blotting using the indicated antibodies. E. Similar experiment as in B, except that U2OS cells were transfected as in D. F. Quantitation of the experiment performed in E. G. SGs assembly was monitored by video-microscopy in parental or H6 NEDP1 KO U2OS cells stably expressing GFP-G3BP1 treated with 0.2mM As (image after 1hr of treatment, video S1A, B). Scale bar: 10μm. Scatter plot showing the average of SGs size (H) and the number of SGs (I) in 50 cells randomly selected per condition (mean ± SD). J, K. Experiment as in G, except that H6 cells were pre-treated with 0.5μM NAEi for 4hrs before treatment with 0.2mM As for 1hr. Scatter plots showing the average of SGs size and number in 50 randomly selected cells from 3 independent experiments per condition (mean ± SD).

Based on these data we also determined the role of NEDP1 in the disassembly of SGs during the recovery period. Alleviation of stress in control cells causes a progressive decrease in the levels of NEDD8 conjugates induced upon stress, indicative of the recovery process (Figure 2A). Under these conditions, using either the GFP-G3BP1 stable cells or endogenous TIA1 as SG markers, we found that compared to control cells, NEDP1 knockout accelerates the elimination of the induced SGs (Figure 2B-E). Similar results were obtained upon transient inhibition of NEDP1 activity by the expression of Nb9 (Figure 2F). The rate of SGs elimination was reduced upon inhibition of the NEDD8 pathway both in control and in NEDP1 knockout cells (Figure 2G). Collectively, these data suggest that: 1. Protein NEDDylation promotes SGs elimination and 2. The acceleration of SGs elimination upon NEDP1 inhibition is due to the increase in protein NEDDylation.

**Figure 2.**
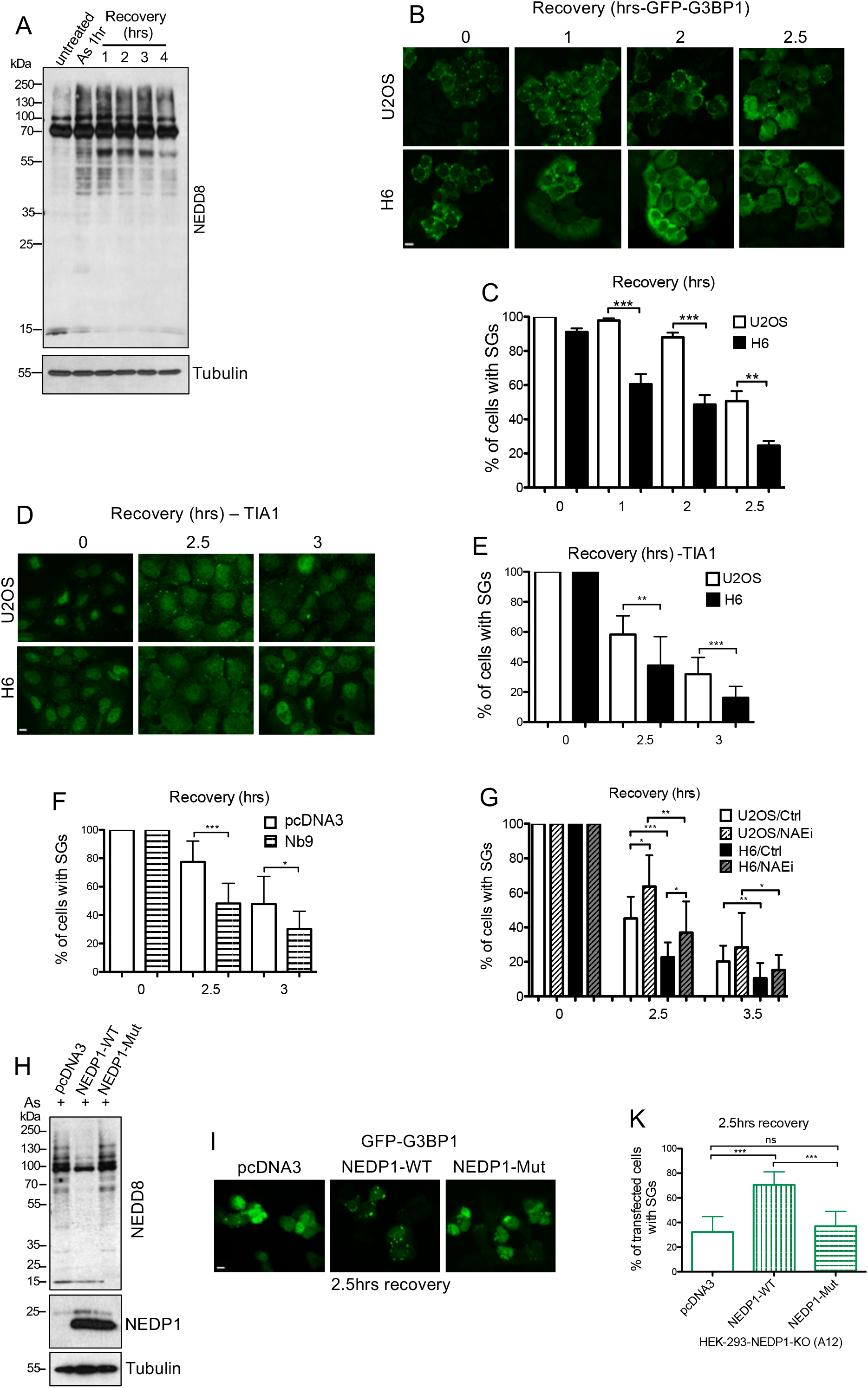
A. U2OS cells were treated with As as indicated and then allowed to recover for the indicated period. Extracts were analysed by western blotting using the indicated antibodies. B. Parental and H6 cells stably expressing GFP-G3BP1 were treated with As (0.2mM) for 1hr before allowed to recover for the indicated periods and fixed. Scale bars, 10μm. C. Quantification of the experiment in B (mean of 3 independent experiments ± SD). D. Similar experiment as in B, using instead parental and H6 U2OS cells and immunostained for endogenous TIA1. E. Quantification of the experiment in D. F. Experiment as described in B, except that cells were prior transfected (48hrs) with empty (pcDNA3) and Nb9 expressing constructs. GFP-G3BP1 staining was used to quantify cells with SGs during the recovery period. G. Parental and H6 U2OS cells stably expressing GFP-G3BP1 pre-treated for 4hrs with 0.5μM NAEi were then treated for 1hr with As (0.2mM), washed with PBS and then incubated in fresh medium containing 0.5μM NAEi for the indicated period before fixation. Graph represents the percentage of cells with SGs (mean ± SD, n=3). H. NEDP1 knockout HEK293 cells (A12) were transfected with the indicated plasmids and 48hrs post-transfection were treated with As (0.2mM) for 1hr before cell extracts were analysed by western blotting with the indicated antibodies. I. NEDP1 KO HEK293 cells (A12) co-transfected with pcDNA3 or NEDP1-WT or NEDP1-Mutant (C163A) and GFP-G3BP1 were treated for 1hr with As (0.2mM) and then allowed to recover for 2.5hrs before fixation and imaging. K. Quantitation of experiment performed in I for percentage of cells with SGs (mean ± SD, n=3).

Next, to determine whether the role of NEDP1 in SG elimination depends on its catalytic activity, we expressed either wild type NEDP1 or its Cysteine catalytic mutant (C163A) in HEK293 NEDP1 knockout A12 cells. As expected, expression of wild type but not of the catalytic NEDP1 mutant decreased protein NEDDylation (Figure 2H). This was accompanied by a reduction in the rate of SG disassembly in wild type NEDP1 overexpressing cells, compared to control or NEDP1 mutant transfected cells (Figure 2I, K). Collectively, the experiments show that inhibition of NEDP1 activity accelerates both the formation and elimination of SGs due to the increase in protein NEDDylation. The observed phenotypes are also not due to defects in microtubule dynamics, which similarly to NEDP1 inhibition result in the formation of smaller and more numerous SGs (Nadezhdina et al., 2010; Wang et al., 2020) (S3, video S2A, B).

### NEDP1 deletion increases the mobility of SG proteins and compromises stress-induced toxicity

A biophysical property that defines the dynamicity and is directly related to the potential pathogenicity of SGs, is the mobility of SG proteins within the condensates. Prolonged stress conditions that can cause extensive protein misfolding reduce the mobility of proteins within SGs (Mateju et al., 2017). The consequence is the transition of SGs from a Liquid-Liquid into a Liquid-Solid phase, where protein-protein interactions prevail on RNA-RNA and RNA-protein interactions. This compromises SG elimination during the recovery period, making these aberrant SGs more dependent on an efficient PQC system for their removal (Ganassi et al., 2016; Mateju et al., 2017). As already discussed, the persistence of such immobile inclusions is regarded as a cause of toxicity (Zhang et al., 2019). Consistent with the above hypothesis, we found by Fluorescence Recovery After Photobleaching (FRAP), that prolonged exposure of parental control cells to As reduced the mobility of GFP-G3BP1 protein with concomitant increase in the half-life of recovery (Figure 3A, B). In contrast, in NEDP1 knockout cells both parameters were unaffected by As (Figure 3A, B), suggesting that NEDP1 inhibition prevents the transition of SGs into a Liquid-Solid phase by maintaining the mobility of SG proteins. Additionally, in survival assays we found that NEDP1 deletion also protects cells against the induced toxicity by As (Figure 3C-E). The data suggest that NEDP1 inhibition promotes SG elimination, which is related to protection of cells against proteotoxicity.

**Figure 3.**
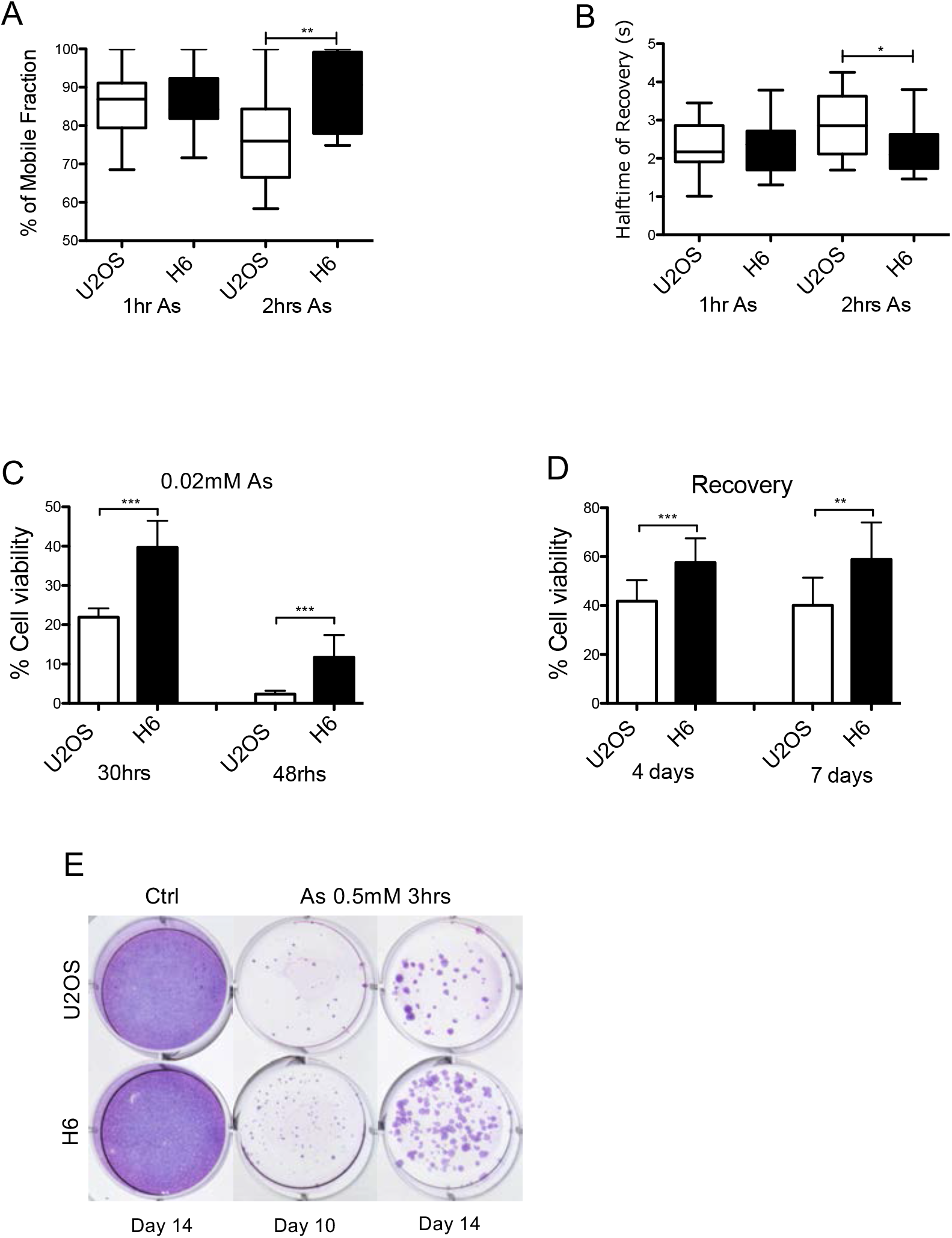
Parental or H6 NEDP1 KO U2OS stably expressing GFP-G3BP1 cells were treated with 0.2mM As and placed on a heated chamber at 37°C with 5% CO2 for the FRAP experiment as described in Star Methods. The Mobile Fraction (A) and Halftime recovery (B) are presented for the indicated period of As treatment. C. Control and NEDP1 KO U2OS cells were plated in 96-well plates and treated continuously with 0.02mM As for 30hrs or 48hrs before viability was measured with the CellTiter-Glo assay. D. Experiment as in C, except cells were treated with As (0.5mM) for 1hr and allowed to recover for the indicated period where cell viability was measured. Graphs represent the mean of the percentage in survival ± SD from 5 independent experiments. E. Parental and H6 U2OS cells were either untreated or treated with As. Clonogenic assay at the indicated periods was performed as described in Star Methods.

### NEDP1 controls SG dynamics through NEDDylation of PARP-1

In order to reveal potential mechanisms for the role of NEDP1 in the control of SG dynamics, we sought to identify NEDP1 substrates. Previous proteomic studies identified hundreds of potential substrates for NEDP1; however these results were based on the use of a NEDD8 mutant (R74K), which may alter the NEDDylome compared to wild type NEDD8 (Lobato-Gil et al., 2020; Vogl et al., 2020). We developed a strategy that allows the identification of NEDP1 dependent NEDDylation sites under endogenous expression of wild type NEDD8. We combined the use of anti-diGly antibodies that recognise both Ubiquitin and NEDD8 modified peptides upon trypsin digestion (Ordureau et al., 2015; Wagner et al., 2011; Xu et al., 2010) with short treatment of cells with the Ubiquitin E1 inhibitor MLN7243 (UAEi), that dramatically reduces Ubiquitin but not NEDD8 modification (S4). We hypothesised that by eliminating the vast majority of Ubiquitin-derived diGly peptides upon MLN7243 treatment, we would be able to quantify diGly peptides derived from NEDD8 modification that depend on NEDP1 (Figure 4A). We previously established the HCT116 colorectal cells as model system for the identification of NEDDylation sites using the diGly approach (Lobato-Gil et al., 2020). We treated parental and NEDP1 knockout cells HCT116 cells (C8) with MLN7243 (UAEi) for 5hrs before extracts were used for trypsin digestion, isolation of diGly modified peptides and mass spectrometry analysis (Figure 4A). We quantified changes in the abundance of 2897 unique modified peptides on 1206 unique proteins between control and NEDP1 knockout cells, and found 934 peptides from 568 proteins and 424 peptides from 328 proteins for which at least a 2-fold increase or decrease respectively was detected upon NEDP1 deletion (Table S1). Our attention was focussed on the targets for which NEDDylation was increased upon NEDP1 deletion, as this would indicate that those proteins are direct NEDP1 substrates. Amongst these targets, we focussed on Poly-(ADP-ribose) polymerase 1 (PARP-1) for several reasons: 1. PARP-1 is a critical regulator of SG dynamics, as PARylation, including of several SGs proteins, generates a scaffold for protein-protein and RNA interactions that is required for SG assembly (Jin et al., 2021). 2. Previous studies indicated that NEDP1 regulates PARP-1 activity upon oxidative stress (Keuss et al., 2019) and 3. Several NEDDylation sites on PARP-1 were also reported in previous proteomic studies (Figure 4B)(Lobato-Gil et al., 2020; Vogl et al., 2020).

**Figure 4.**
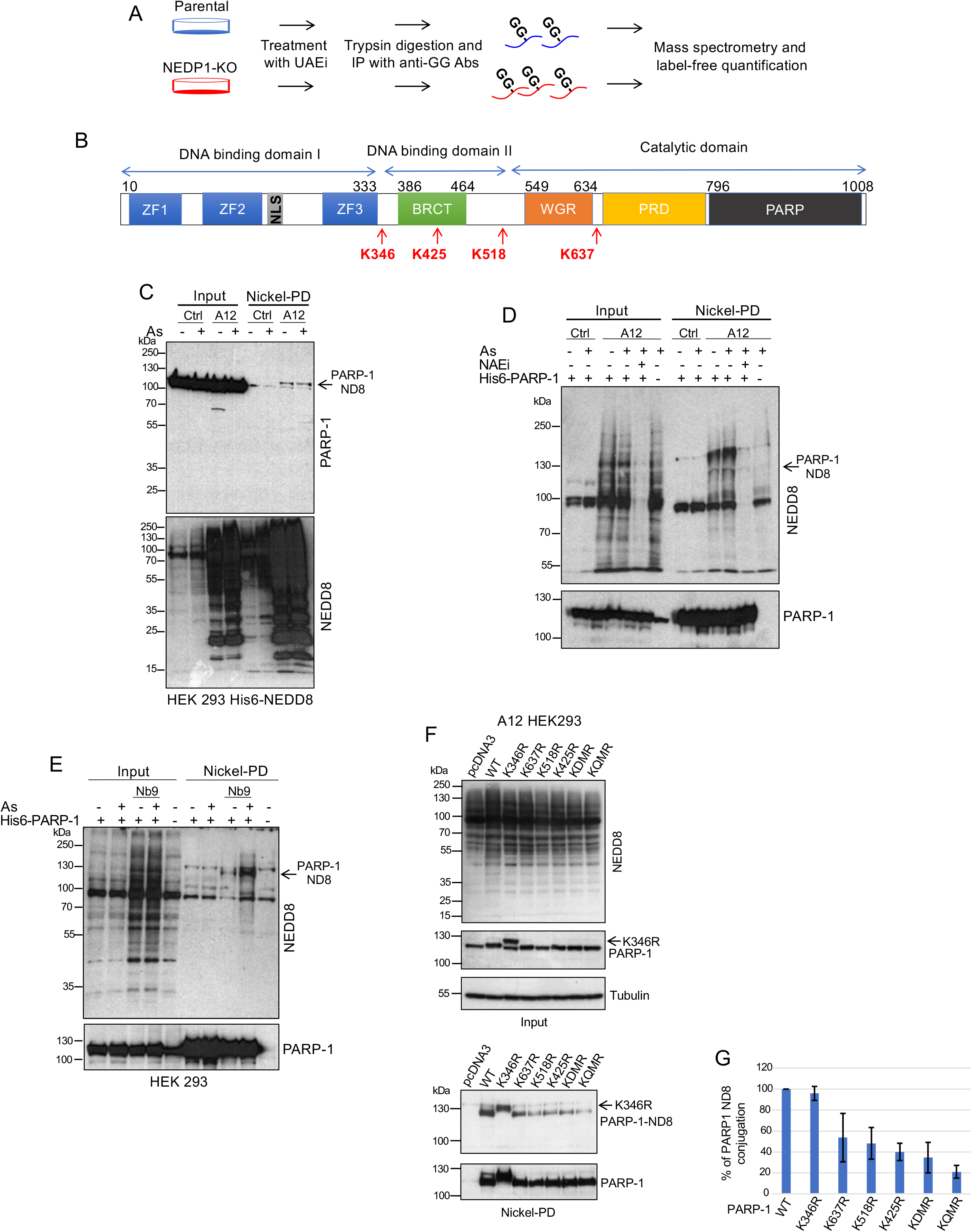
A. Schematic representation of the used proteomic approach to identify NEDDylation sites controlled by NEDP1 at endogenous levels of wild type NEDD8 expression. Parental and NEDP1 knockout (C8) HCT116 cells were treated with UAEi (0.5*μ*M, 5hrs) before extracts were used for trypsin digestion and immunoprecipitations with anti-diglycine antibodies and mass spectrometry analysis. Data were used for label-free quantification. B. Schematic representation of PARP-1 domains including the identified NEDDylated lysine residues upon NEDP1 knockout (this study, Vogl et al., 2020; Lobato et al., 2021). C. Parental and NEDP1 knockout (A12) HEK293 cells stably expressing His6-NEDD8 were treated with As (0.5mM, 1hr) before extracts were used for Nickel-PD. Isolated His6-NEDD8 conjugates and total cell extracts (input) were used for western blot analysis with the indicated antibodies. D. Parental and A12 cells were transfected with His6-PARP-1. 48hrs post-transfection cells were treated with As (0.5mM, 1hr) as indicated and extracts were used for Nickel-PD and western blot analysis as in C. E. HEK293 cells were transfected His6-PARP-1 and Nb9 expression constructs as indicated. 48hrs post-transfection, cells were treated with As (0.5mM, 1hr) and Nickel-PD and western blot analysis was performed as in D. F. A12 HEK293 cells were transfected with the indicated His6-PARP-1 expression constructs and 48hrs later were used for Nickel-PD. Isolated proteins and total cell extracts (input) were used for western blot analysis as indicated. G. Quantitation of the experiment performed in F. Values represent the average of 3 independent experiments ± SD.

We first determined if PARP-1 is a NEDP1 dependent NEDD8 substrate. We isolated NEDD8 conjugates from control and NEDP1 knockout HEK293 cells (A12) stably expressing His6-NEDD8 and monitored by Nickel pull-down (Nickel-PD) and western blotting the NEDDylation of endogenous PARP-1 (Figure 4C). Deletion of NEDP1 (A12) or inhibition with Nb9 dramatically increases PARP-1 NEDDylation compared to control cells, consistent with the proteomics studies (Figure 4C-E). Exposure of cells to As did not have a significant effect on PARP-1 NEDDylation (Figure 4C, D), which was however dramatically reduced upon treatment with the NAE inhibitor MLN4924 (Figure 4C, D, NAEi). This shows that NEDP1 dependent PARP-1 NEDDylation depends on the canonical NEDD8 pathway (Meszka et al., 2022). To characterise the role of PARP-1 NEDDylation in SGs dynamics we generated a series of PARP-1 mutations in Lysine residues (K364, K425, K518, K637) identified as potential NEDDylation sites in current and previous proteomic studies (Figure 4B, F, Table S1) (Lobato-Gil et al., 2020; Vogl et al., 2020). Wild type and mutant His6-PARP-1 constructs were expressed in control and NEDP1 knockout (A12) HEK293 cells and their modification with endogenous NEDD8 was monitored upon isolation of His6-PARP-1 by Nickel-PD (Figure 4F). We found that mutation of K425, K518, K637 into Arginine or the generation of a double mutant (K518/637R, KDMR) significantly but not completely reduced PARP-1 NEDDylation (Figure 4F, G). No effect was observed upon mutation in K346, which intriguingly altered the migration profile of PARP-1 (Figure 4F, G). Mutation of all four Lysines into Arginine (quadruple mutant, KQMR) caused the most profound reduction in PARP-1 NEDDylation (Figure 4F, G). This analysis indicates that multiple sites in PARP-1 are used for NEDDylation, with K425 and K518 as the most prominent.

To assess the potential role of PARP-1 NEDDylation in SG dynamics, we used the KQMR and the single K425R mutant as they displayed the most reproducible reduction in PARP-1 NEDDylation (Figure 4G). To avoid any interference from the expression of endogenous PARP-1, we used U2OS cells where PARP-1 has been knocked out (PARP-1^-/-^) (Hanzlikova et al., 2016) and we also generated double PARP-1^-/-^ / NEDP1^-/-^ cells by CRISPR/Cas9 (Bailly et al., 2019). Expression of either of the two NEDDylation defective PARP-1 mutants, K425R, KQMR in these model cell lines results in higher levels of PAR upon As treatment compared to wild type PARP-1, indicating that NEDDylation compromises PARP-1 activity (Figure 5A). Consistent with this notion, we found that in NEDP1 knockout cells with increased levels of PARP-1 NEDDylation (Figure 4C, D), the levels of PAR induced upon As are lower compared to control parental cells (Figure 5B). By monitoring SG dynamics using endogenous TIA1 as SG marker, we found that compared to the expression of wild type PARP-1, the expression of the NEDDylation defective PARP-1 mutants is corelated with an increase in the size of SGs, especially during the recovery period (Figure 5C, D). In addition, SG disassembly was compromised in cells expressing the NEDDylation deficient PARP-1 mutants, compared to cells expressing wild type PARP-1 (Figure 5E). Collectively, the data support the following hypothesis: NEDP1 inhibition causes the hyper-NEDDylation of PARP-1, reducing the production of PAR. This results in the formation of smaller SGs, which are eliminated faster during the recovery period compared to control cells (S5A).

**Figure 5.**
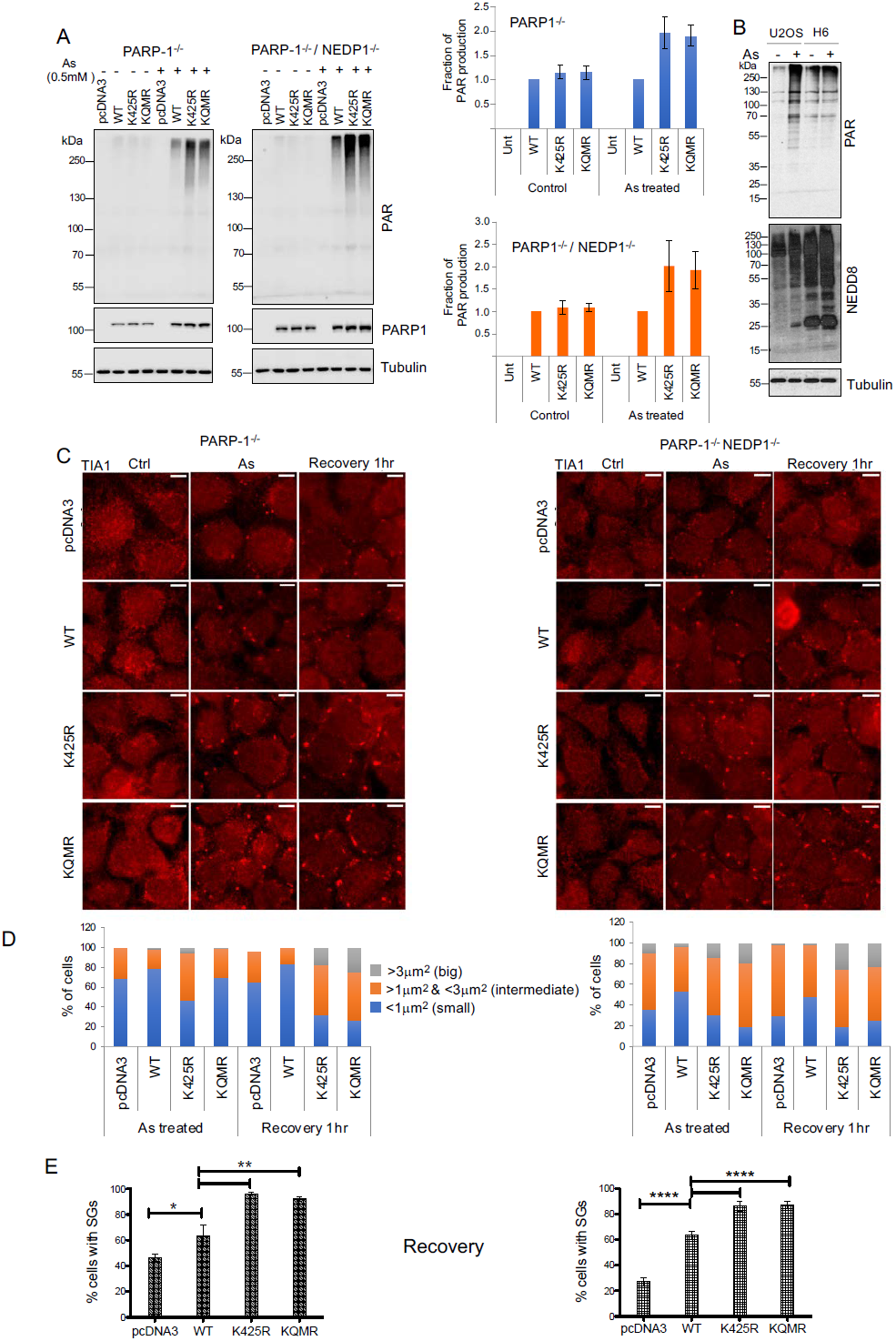
A. (Left panels). PARP-1^-/-^ and PARP-1^-/-^ / NEDP1^-/-^ U2OS cells were transfected with the indicated plasmids expressing WT or NEDDylation deficient PARP-1 mutants. 48hrs post-transfection cells were either untreated or treated with As (0.5mM, 1hr) and total cell extracts were used for western blot analysis with the indicated antibodies. (Right panels). Quantitation of the experiment on the left panel. Values represent the mean ± SD from 5 independent experiments of the fraction of PAR production (WT was used as the reference for each condition). B. Parental or H6 U2OS cells treated with As (0.5mM, 1hr) and extracts were used for western blot analysis with the indicated antibodies. C. Experiment was performed as described in A, except that recovery was also employed as indicated. Formation of SGs was monitored by immunostaining for endogenous TIA1. D. In the experiment performed in C, the size of SGs was measured. Values represent the mean of the percentage of cells with the indicated size of granules. E. Quantitation of the experiment performed in C, showing the percentage of cells with SGs during the recovery period. Values represent the mean ± SD from 3 independent experiments.

To further assess the significance of the effect of NEDP1 in SGs dynamics through PARP-1 activity regulation, we performed an “ epistasis-like” experiment. We monitored the effect of PARP-1 inhibition on SG disassembly in control and NEDP1 knockout cells. Consistent with the established function of PARP-1 in SG dynamics, chemical inhibition of PARP-1 with Olaparib, accelerates the elimination of SG in control cells. However, no further acceleration was observed in NEDP1 knockout cells (S5B). This is consistent with the idea that NEDP1 and PARP-1 are functionally linked and NEDP1 inhibition accelerates SG elimination mainly through hyper-NEDDylation of PARP-1, compromising the production of PAR.

### NEDP1 inhibition accelerates the elimination of ALS-related SGs

Mutations in many SGs-related proteins identified in neurodegenerative diseases such as ALS/FTD, are regarded as the cause of formation of pathogenic SGs. A key characteristic of such inclusions is their persistence in the cytoplasm compared to the relative fast elimination of physiological SGs during recovery. We tested the effect of NEDP1 inhibition on the elimination of SGs containing well-established ALS-derived mutants for TIA1 (A381T and E384K) (Mackenzie et al., 2017). As expected, elimination of SGs containing mutant TIA1 was slower compared to SGs containing wild type TIA1 in control cells (Figure 6A, B). Interestingly, NEDP1 knockout accelerated the elimination both of wild type and mutant TIA1 SGs to almost the same extent (Figure 6A, B). Employing similar assays, we found that NEDP1 activity inhibition by Nb9 significantly accelerated the elimination both of wild type and mutant TIA1 SGs compared to control transfected cells (Figure 6C). We next assessed whether NEDP1 inhibition promotes the elimination of aberrant SGs using more physiologically relevant *in vitro* systems. To this end, we performed similar experiments in mouse-derived hippocampal neurons. Transfection of hippocampal neurons with Nb9 accelerated the elimination of both wild type and mutant GFP-TIA1 SGs, similarly to what is observed in transformed cells (Figure 6D, E). Additionally, we determined the effect of NEDP1 inhibition on SG elimination in skin fibroblasts derived from a patient affected by sporadic ALS (sALS 37/15). To achieve high level of NEDP1 inhibition in these cells, we employed infection with retrovirus expressing Nb9. Compared to fibroblasts from healthy donors (HDF20), the elimination of SGs (eIF4G1 staining) during the recovery period is significantly delayed in sALS 37/15 fibroblasts, consistent with the correlation between persistence of aberrant SGs and ALS pathology (Figure 6F). Infection with a retrovirus expressing Nb9, accelerated the elimination of SGs both in healthy HDF20 and sALS 37/15 fibroblasts compared to control infected cells (Figure 6F). Using an empty viral vector as a control, we excluded that viral infection per se could influence SGs dynamics (S6). Collectively, these data reveal that NEDP1 inhibition accelerates the clearance of pathogenic-related SGs in non-neuronal immortalized cell lines, in primary neuronal cells, as well as in patient-derived fibroblasts.

**Figure 6.**
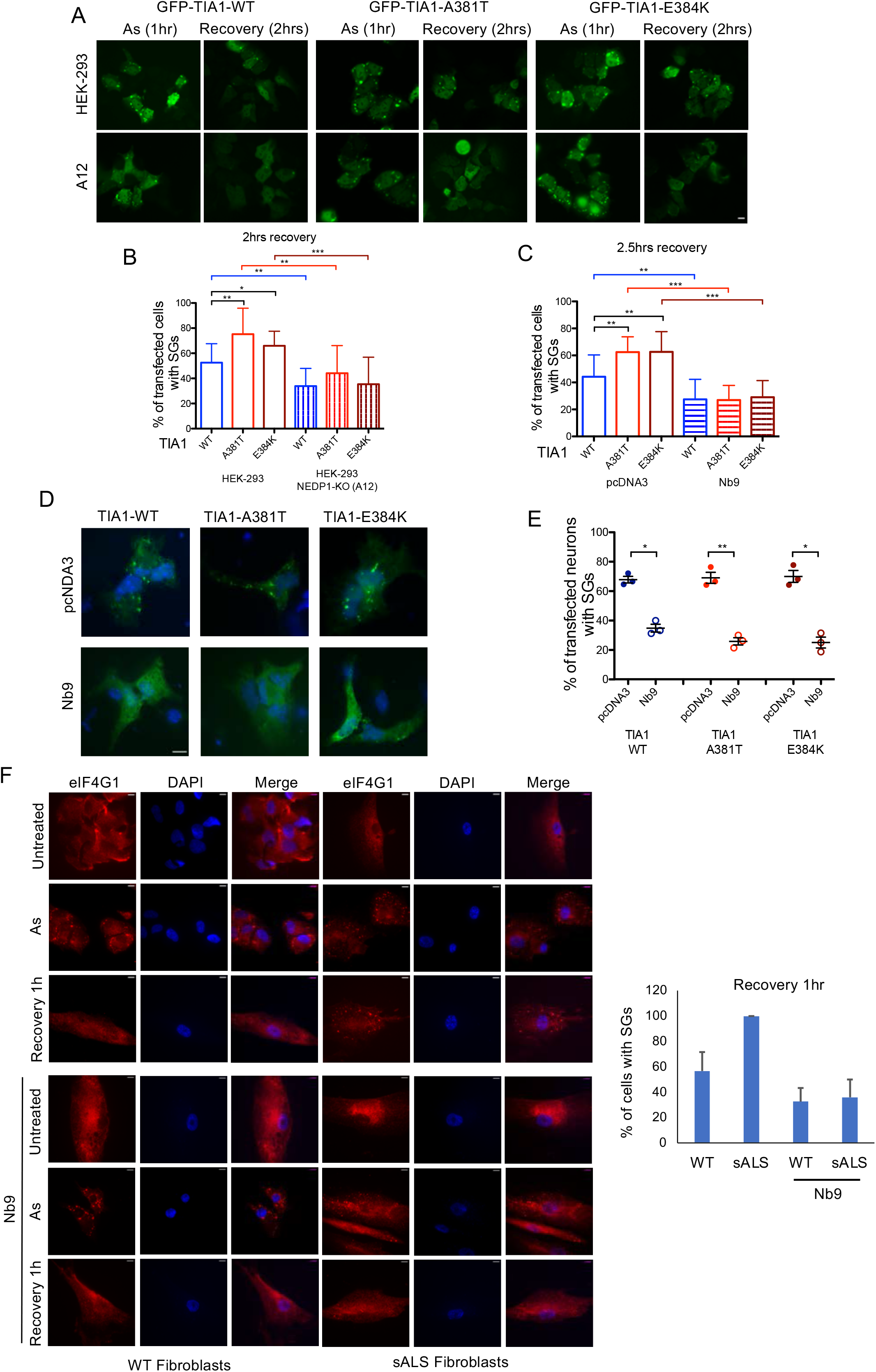
A. Parental or A12 (NEDP1 knockout) HEK293 cells were transfected with the indicated GFP-TIA1 constructs (1*μ*g) and 48hrs post-transfection cells were stressed with As (0.5mM, 1hr) before recovery (2hrs). SGs formation was monitored by GFP fluorescence. B. Quantitation of the experiment performed in A. Values represent the mean of 3 independent experiments ± SD for the percentage of cells with SGs. C. Experiment performed as in A, except that where indicated the Nb9 was co-transfected. Quantitation was performed as in B. D. Mouse-derived hippocampal neurons were transfected with the indicated GFP-TIA1 and Nb9 constructs, stressed with As (0.2mM, 1hr) before allowed to recover for 2hrs. SGs formation was monitored by GFP fluorescence. E. Scatterplot representing the percentage of co-transfected cells with SGs (mean ± SD). Each dot represents the percentage of co-transfected neurons with SGs on one coverslip (3 coverslips with 20-25 co-transfected neurons per condition). F. Human fibroblasts derived from healthy (WT) donors or patients with sporadic ALS (sALS) were infected with retrovirus expressing NB9 as indicated. Cells were either untreated or treated with As (0.5mM, 1hr) and allowed to recover (1hr). SG formation was monitored by immunostaining for the SG protein eIF4G1. Right panel represents the quantification of the experiment. Values represent the mean ± SD from 3 independent experiments of the percentage of cells with SGs.

### NEDP1 deletion causes the elimination of aberrant SGs and restores motility in a *C. elegans* ALS model

To assess the role of NEDP1 on SG formation at the organism level, we used the worm *C. elegans* as model system. *C. elegans* is an established model for neurodegenerative diseases, including ALS. A well-established *C. elegans* ALS model is based on the introduction of human ALS-linked SOD1 mutations in the worm homologous gene *sod-1* (Baskoylu et al., 2018). Expression of SOD1 mutants such as G85R in *C. elegans*, leads to a strong motility defect associated with the presence of insoluble aggregates of the SOD1 G85R protein (Forsberg et al., 2010), recapitulating an ALS-related cellular pathology.

SG formation can be monitored in living animals by the expression of an endogenous GFP-tagged version of the G3BP1 worm homologous gene *gtbp-1* (Paix et al., 2014). We have previously defined the worm *ulp-3* as the homologous gene to human NEDP1 and generated animals deleted for *ulp-3* (Bailly et al., 2019). Our studies on these animals identified a cytoplasmic role for ULP-3 on apoptosis execution in response to genotoxic stress that is highly conserved in human cells (Bailly et al., 2019).

Taking advantage of this knowledge and tools, we studied the role of ULP3 in SG formation in *C. elegans*. We first monitored SG formation in response to proteotoxic stress induced either by heat shock or As, similarly to the performed assays in human cells. Exposure of wild type *gtbp-1::gfp* animals to both stress conditions induced the formation of GTBP-1 protein puncta, indicative of SG formation in muscle pharyngeal cells, as previously reported (Figure 7A, B) (Kuo et al., 2020). In contrast, in *gtbp-1::gfp* animals deleted for *ulp-3*, the number of detectable GTBP-1 SGs was dramatically reduced under both stress conditions (Figure 7A, B).

**Figure 7.**
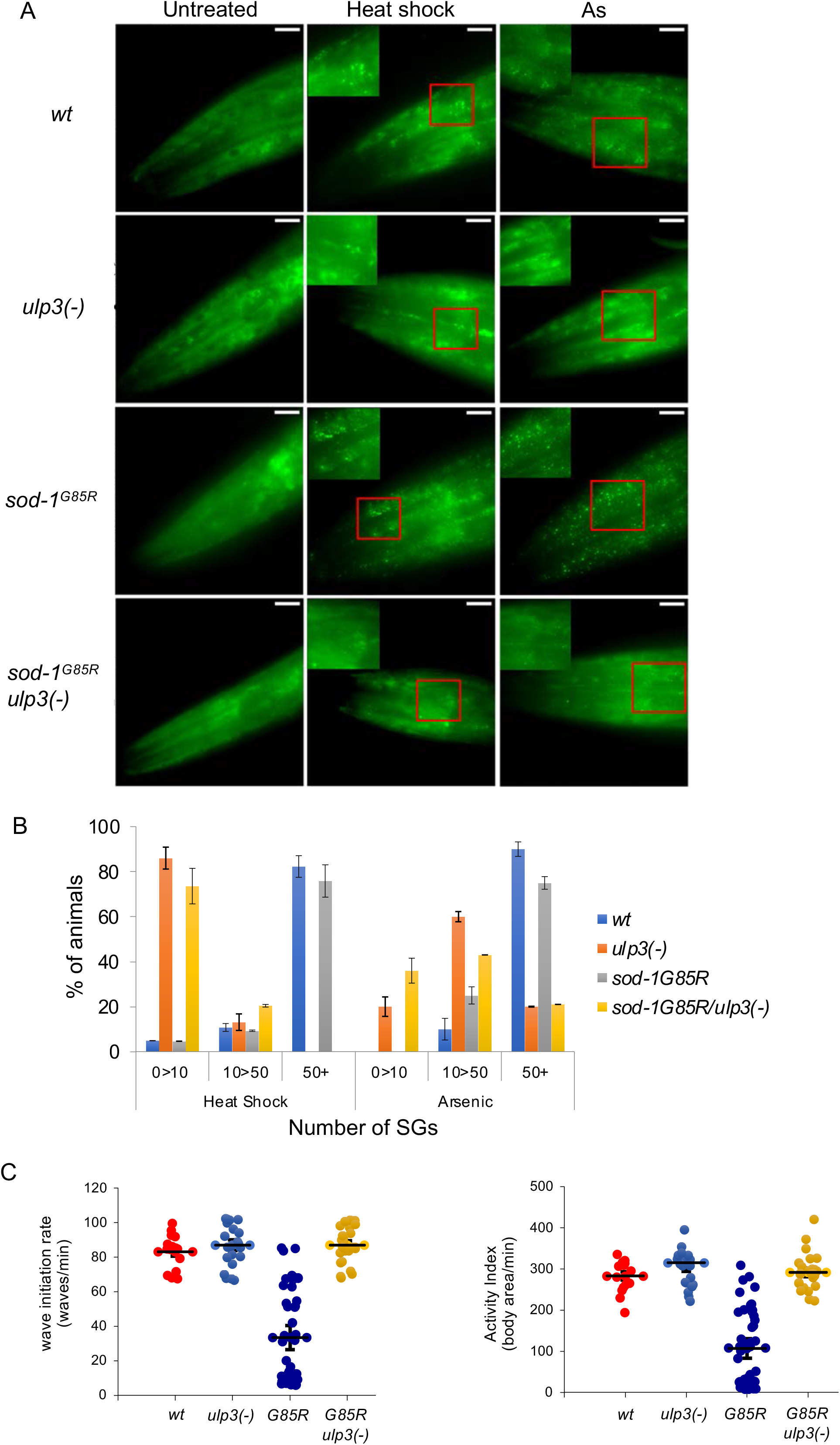
A. Animals expressing GFP-GTBP-1 on the indicated genetic background where either untreated or exposed to heat shock or As as described in Star Methods. The formation of SGs was monitored by GFP fluorescence. B. Quantitation of the experiment performed in A. Values are the mean of 2 independent experiments ± SD of the percentage of animals (n>10/experiment) with the indicated number of SGs. C. Motility experiments were performed as described in Star Methods using 2 different parameters, wave initiation rate (left) and activity index (right).

To determine the role of ULP3 in ALS-pathology we monitored SG formation in *gtbp-1::gfp* animals expressing the G85R mutated form of SOD1. In particular, As treatment generated larger GTBP-1-GFP SGs compared to control animals (S7), consistent with the formation of aberrant inclusions in the muscle pharyngeal cells and presumably in the surrounding neurons (Figure 7A, B). Strikingly, under both stress conditions, deletion of *ulp-3* dramatically reduced the formation of GTBP-1-GFP SGs in the *sod-1*^G85R^ mutant background (Figure 7A, B). Collectively, the analysis in human cells and in *C. elegans* revealed a highly conserved role for NEDP1/ULP3 in the elimination of physiological and pathological ALS-related SGs. Considering that oxidative stress aggravated the size of SGs specifically in the single copy *sod-1*^G85R^ mutation carrying animals, we examined the effect of oxidative stress on worm locomotion. We used an established method, where animals in liquid culture are exposed to oxidative stress (paraquat) overnight and the quantitative analysis of their swimming behaviour, indicative of animal motility, is performed using the software CeleST (Restif et al., 2014). By measuring two different parameters of animal motility (Wave Initiation Rate and the Activity Index), we found that the *sod-1*^G85R^ mutant showed a severe defect in locomotion compared to the wild type animals (Figure 7C, video S3A, B). Critically, *ulp-3* deletion had a dramatic effect on *sod-1*^G85R^ animal locomotion, as it restored the observed phenotypic defects almost back to wild-type levels (Figure 7C, video S3C, D). Overall, our results establish a correlation between the accumulation of ALS-related SGs and locomotion defects; these data are in line with previous findings in cells showing that the conversion of persisting SGs into pathological aggregates causes cell toxicity (Zhang et al., 2019). Importantly, our data suggest that the elimination of ALS-related SGs observed in the *sod-1*^G85R^ mutant upon *ulp-3* deletion is correlated with the rescue of ALS-phenotypes related to animal motility.

## Discussion

### NEDP1 controls SG dynamics through PARP-1 NEDDylation

The formation of SGs is a generalized cellular response, regarded as a protective mechanism to a broad range of stress stimuli. The assembly and disassembly processes collectively characterize the dynamic nature of SGs, which is the essence of their protective role in stress responses. Mutations found in neurodegenerative diseases such as ALS contribute to the formation of aberrant SGs that can mature into coalesce with pathological inclusions (Zhang et al., 2019). One widely accepted hypothesis, is that the persistence of such inclusions, is a molecular basis for the progression of the disease. Thus, developing approaches that restore SG disassembly and avoid their conversion into an aggregated state would contribute to eliminate such toxic inclusions and represents an attractive approach for therapeutic intervention.

Here, we identify the deNEDDylating enzyme NEDP1 as regulator of SGs dynamics. Using multiple approaches, including the use of a Nb as first-in class inhibitor for NEDP1, we found that NEDP1 inhibition promotes the elimination of both physiological and aberrant SGs found in ALS cell and animal models, including patients’ derived fibroblasts. As NEDP1 is a highly specific component of the NEDD8 cycle, the study provides strong evidence for the role of the NEDD8 pathway as regulatory component of SG dynamics.

Detailed kinetic analysis revealed an intriguing role for NEDP1 in SGs dynamics, as its inhibition accelerated both the assembly and disassembly process. The formation and dissolution of SGs are regarded as multi-step events. The initial formation of the core is followed by the progressive recruitment of the shell proteins and/or by fusion events that result in the growth of SGs (Jain et al., 2016; Wheeler et al., 2016). Whether these different steps in SG assembly follow the same rates is not clear. Based on the observation that NEDP1 knockout cells produce higher number but smaller SGs compared to wild type cells, the apparent acceleration in SG assembly may reflect a defect in the subsequent growth/fusion of SGs. This in turn, may also be the basis for the accelerated SG dissolution observed during the recovery process. As NEDP1 also controls the dynamics of pathological aberrant SGs, the above-described characteristics reveal a fundamental function of NEDP1 in SG biology.

Mechanistically, we identified PARP-1 as the NEDD8 substrate through which NEDP1 controls SG dynamics. Several lines of evidence have established that the PARP-1 product, PAR, either by directly modifying substrates or through formation of unanchored chains creates the scaffold that allows the formation of SGs (Grimaldi et al., 2019; Jin et al., 2021). Whether PAR equally affects the different steps in SG formation (core/shell formation, fusion) is still unclear. What is however emerging is that PAR production has to be modulated, especially for SG disassembly. Excessive production of PAR, due to PARP overactivation and/or inhibition of PAR glycohydrolases (PARGs), delays SG dissolution during recovery and results in the formation of aberrant inclusions especially for disease-related mutant SG proteins (Duan et al., 2019; Grimaldi et al., 2019; Jin et al., 2021). The hyper-NEDDylation of PARP-1 upon NEDP1 inhibition compromises PAR production, thereby promoting SG disassembly, as demonstrated by the mutational analysis on NEDDylation deficient PARP-1 mutants. Consistent with the above-described functional link between NEDP1 and PARP-1, inhibitors for PARP-1 had no additive/synergistic effects on SGs dissolution in NEDP1 knockout cells. Collectively, the data indicate that NEDP1 may indeed play a modulatory role for PARP-1 activity to finely control PAR production and SG dynamics during the stress response. This notion is further supported from the observation that protein NEDDylation is increased upon oxidative stress (Figure 1A) to potentially “ buffer” PARP-1 activity and subsequently modulate SG dynamics.

### NEDP1 as potential therapeutic target in ALS

Mutations in several genes including SOD1, TIA1, FUS, TDP-43 have been implicated in the generation of aberrant protein inclusions with impaired kinetics that contain several SG proteins and are prominent in ALS patients (Bosco et al., 2010; Dewey et al., 2011; Lee et al., 2020; Patel et al., 2015). The persistence of such aberrant inclusions due to defects in their elimination is regarded as a molecular basis of the disease.

The combination of studies in human cells, mouse neurons, ALS-patient fibroblasts and in *C. elegans* as model organism for ALS, show that NEDP1 inhibition can accelerate the elimination of pathologic aberrant SGs regarded as hallmark in ALS. Critically, the data in *C. elegans* demonstrate that the elimination of ALS-related SGs upon NEDP1 inhibition is correlated with restoration of animal motility, providing the proof-of-principle for targeting NEDP1 for therapeutic intervention. This notion is strongly supported with the use of Nbs that potently and specifically inhibit NEDP1 activity (Abidi et al., 2020) and fully reproduce the findings on aberrant SG dissolution using genetic approaches for NEDP1 inhibition.

Based on the established role of PARP-1 in SG dynamics, PAPR-1 inhibitors have been proposed as an attractive therapeutic approach for the elimination of aberrant SGs. A complication with this approach is that in addition to the elimination of pathogenic inclusions, PARP-1 inhibitors also induce DNA damage and cell toxicity, effects that are desired for cancer treatment but not for the treatment of neurodegenerative diseases (Puentes et al., 2021). A proposed concept to overcome this problem, is the development of low toxicity PARP-1 inhibitors that will not impair the DNA damage response, but will still retain their ability to prevent PARP-1 hyperactivation and promote aberrant inclusion elimination (Puentes et al., 2021). NEDP1 knockout/inhibition in several organisms including *C. elegans, Drosophila, Arabidopsis, S. pombe* has no effect on survival/development, but rather plays an important role in the cellular stress responses (Meszka et al., 2022). In particular, genetic studies showed that NEDP1 inhibition blocks apoptosis induction in response to DNA damage and improves cell survival, with no effect in the DNA damage checkpoint signalling pathway (Bailly et al., 2019). Thus, it is predicted that targeting NEDP1 will provide the desired modulation of PARP-1: It will compromise PARP-1 hyperactivation and promote the elimination of pathogenic inclusions, in the absence of any toxic effects.

The study reveals NEDP1 as a potential target to ameliorate ALS-related phenotypes and provides the proof-of-principle for the use of the developed anti-NEDP1 Nbs as attractive therapeutic agents for ALS.

## Acknowledgements

The study was financially supported by the Labex EpiGenMed, an " Investissements d’avenir " program, reference ANR-10-LABX-12-0, a postdoctoral fellowship to RS by FRM (SPF202110013944) and the European commission, Horizon 2020 that supports the UbiCODE Research Training network. We are grateful to the imaging facility MRI, member of the national infrastructure France-BioImaging infrastructure supported by the French National Research Agency (ANR-10-INBS-04, ≪Investments for the future≫)”. We also thank Maxime Bonnet (Anne Debant’s laboratory, CRBM), for providing mouse-derived hippocampal neurons and Benjamin Vitre (Benedicte Delaval’s laboratory, CRBM) for the EB1-mcherry expressing retrovirus. SC and JM acknowledge AriSLA (Granulopathy and MLOpathy) for the generation and study of human fibroblast lines. Protocols and informed consent for the generation of the HDF20 and sALS 37/15 fibroblast lines were approved by the Institutional Ethics Committee (Protocol n375/04, 07/01/2004 and Protocol n 299/14, 28/04/2015). We thank Dr. Cristina Cereda for the generation of the human fibroblast lines. SC also acknowledges Departments of excellence 2018–2022; E91I18001480001.

## Authors contributions

TK and RS performed the majority of experiments. IM and AB identified and characterised the NEDDylation sites on PARP-1, including the generation of PARP-1 mutants. HT and JP assisted with the SGs dynamics experiments, survival/clonogenic assays and the generation of Nb expressing virus. RS and AB performed all experiments in *C. elegans* as model organism for ALS. JM and SC generated and provided the ALS-fibroblasts. DPX with the help of all members conceived and managed the project. DPX wrote the manuscript with the assistance of SC.

## Supplemental Videos

Video S1. Live imaging of SG dynamics in parental (A) and H6 (B) U2OS cells expressing GFP-G3BP1. Related to Figure 1.

Video S2. Microtubule dynamics in parental (A) and H6 (B) U2OS cells expressing EB1-mcherry. Related to Figure 1.

Video S3. *C. elegans* motility upon exposure to oxidative stress (paraquat). Related to Figure 7.

## Materials and Methods

### Materials

The following antibodies were used: Rabbit monoclonal anti-NEDD8, Y297 (#GTX61205, Abcam), rabbit anti-H2A (#Ab13923, Abcam), rabbit anti-ubiquitin (#z0458, DAKO), mouse anti-PARP-1 (F-2) (#sc-8007, Santa Cruz), mouse anti-TIA-1 (G-3) (#sc-166247, Santa cruz), mouse anti-eIF4G (A-10) (#sc-133155, Santa Cruz), mouse anti-PAR (10HA) (#4335-MC-100, Trevigen), mouse anti-GFP (#11814460001, Roche), mouse anti-tubulin (#3873, Cell Signaling), sheep anti-NEDP1 (in house)(38), mouse anti-Flag (#A2220, Sigma-Aldrich), Ubiquitin Branch Motif Antibody (K-ε-GG), (#3925, Cell Signaling), secondary antibodies (anti-mouse #4416, anti-rabbit #A0545, anti-sheep #A3415, Sigma-Aldrich). MLN4924 (NAEi) was purchased from Active Biochem (#A-1139), MLN7243 (UAEi) from Chemietek (#CT-M7243), Fugene6 HD (#E2691, Promega), lipofectamine 2000 (#11668019, Invitrogen), Ni-NTA Agarose (#30210, QIAGEN), PVDF membrane (#LC2002, Millipore), ECL Western Blotting Detection Reagents (#RPN2232, Amersham), CellTiter-Glo® Luminescent Cell Viability Assay (#G7570, Promega). Methyl metanosulfonate (MMS, #129925), camptothecin (CPT, #C9911) and etoposide (Eto, #E1383) from Sigma-Aldrich.

### Worm strain and culture conditions

Hermaphrodite worms were maintained at 20°C on Nematode Growth Media (NGM) agar plates seeded with E. coli strain OP50. NGM plates contain 1.5% agar, 0.25% Tryptone, 0.3% Sodium Chloride, 1mM Calcium Chloride, 1mM Magnesium Sulfate, 25mM Potassium Phosphate (pH 6.0), 5μg/ml Cholesterol. Double and triple mutants were isolated by PCR and confirmed by genomic sequencing. Mutants generated in this study will be deposited at the CGC and/or will be provided on request.

The following strains were used in the study:

wt: N2 (Bristol)

ulp-3(tm1287) IV (DPX1287)

sod-1(rt449[G85RC]) II (HA2987)

gtbp-1(ax2055[gtbp-1::GFP]) IV (JH3199)

ulp-3(tm1287) IV, gtbp-1(ax2055[gtbp-1::GFP]) IV (DPX11)

sod-1(rt449[G85RC]) II; gtbp-1(ax2055[gtbp-1::GFP]) IV (DPX12)

sod-1(rt449[G85RC]) II; gtbp-1(ax2055[gtbp-1::GFP]) IV, ulp-3(tm1287) IV (DPX13)

csIs(pqn-59::GFP) I; gtbp-1::RFP(ax5000) IV (HML713)

kasls7[snb-1p::C9 ubi+ myo-2::GFP] (KRA315)

kasls7[snb-1p::C9 ubi+ myo-2::GFP]; ulp-3(tm1287) IV (DXP20)

kasls7[snb-1p::C9 ubi+ myo-2::GFP]; gtbp-1::RFP(ax5000) IV (DPX21)

kasls7[snb-1p::C9 ubi+ myo-2::GFP]; gtbp-1::RFP(ax5000), ulp-3(tm1287) IV (DPX22)

gtbp-1::RFP(ax5000) IV (DPX23)

gtbp-1::RFP(ax5000), ulp-3(tm1287) IV (DPX24)

### Fibroblasts

The following set of human-derived fibroblasts were used:

HDF20 Female 63 years-old, fibroblast passage 9

37/15 sALS Female 59 years-old, fibroblast passage 6

### Cell culture

Cell lines, U2OS (Female) preferred system for live imaging, HCT116 (Female) preferred system for diGly proteomics based on previous studies (44), HEK293 (Male), preferred system for transient over-expression studies due to high transfection efficiency were originally obtained from the ATCC bioresource. Cell lines were maintained in DMEM, in 10% FCS and standard antibiotics (streptomycin, penicillin), 5% CO2 and 37oC and regularly tested for mycoplasma contamination. Cell lines have not been authenticated. Stable cell lines expressing His6-NEDD8 where established using puromycin 2.5*μ*g/ml as described in (27). For cell lines stably expressing GFP-G3BP1, cells were selected with G418 (1mg/ml) for 14 days before a pool of cells stably expressing GFP-G3BP1 was acquired. U2OS NEDP1 knockout cells were generated by CRISPR/Cas9 as previously described. Using similar approaches were used to generate the HCT116 and HEK293 NEDP1 knockout cells. The deletion within the NEDP1 gene was confirmed by DNA sequencing and by western blot analysis the absence of NEDP1 protein with concomitant accumulation of NEDD8 conjugates. Parental and NEDP1 knockout U2OS cells were infected with EB1-mcherry expressing vector virus (gift from Dr Benjamin Vitre). Cells were selected with puromycin. Dissected mouse-derived hippocampal neurons were plated in medium A (Neurobasal, B27, 10% foetal calf serum, Glutamax, 25[M glutamic acid, pen/strep) for 2hrs before changing into medium B (Neurobasal, B27, Glutamax, 0.45% glucose, pen/strep), whereas fibroblasts derived from ALS patients were grown in high glucose DMEM (without Sodium Pyruvate), supplemented with 2mM L-glutamine, 100U/ml penicillin/streptomycin and 10% foetal bovine serum.

### Transfection

U2OS cells were transfected with the indicated plasmids with Fugene 6 (Roche) according to manufacturer’s instructions using a 3:1 Fugene/plasmid ratio. For HEK293 the calcium/phosphate method was employed. Transfection of mouse derived hippocampal neurons was performed using lipofectamine 2000.

### Retrovirus production

The Nb9 expressing DNA was cloned in the pMXs-Puro retroviral vector and co-transfected in 10cm cell-culture dishes of 5×10^6^ HEK293T cells, with virus packaging and envelope expressing plasmids using Fugene-6 (Roche). Medium was replaced with DMEM with 10% FCS the next day and supernatant was harvested 3 days post-transfection, filtered through a 0.45*μ*m pore-size filter (Sartorius, Minisart), and stored at −80^°^C.

### Isolation of His6-tagged proteins

Isolation of His6-NEDDylated proteins and His6-PARP-1 was performed under denaturing conditions as described in (28).

### SGs assembly and recovery

For assembly, cells grown on coverslips in a 12-well plate were treated with 0.2 or 0.5mM sodium arsenite for the indicated time before washing 3 times with PBS followed by fixation with 3.7% paraformaldehyde. For recovery, the medium containing arsenite was removed and the cells were washed 3 times with PBS then incubated in fresh medium for the indicated time before fixation. Under the used arsenite treatment (1hr, 0.5mM) almost 100% of cells form SGs in all conditions and this was the point of recovery. Formation and recovery of SGs was evaluated by subsequent immunocytochemistry assays. Images of 15 random fields (more than 100 cells) were selected on each coverslip to quantify the percentage of cells with SGs in each condition. The size and the number of SGs, was manually measured with ImageJ. In the experiments where NAEi and UAEi were used the equivalent volume of DMSO (solvent used to dissolve the compounds) was added in the untreated/control conditions. For hippocampal neurons, cells were seeded in 6-well plates on coverslips coated with poly-L-ornithine (0.25mg/ml). 30min before transfection, fresh medium with no glucose was added and cells were transfected with TIA1 (1*μ*g) and Nb9 (2*μ*g) expression plasmids using lipofectamine 2000. 48hrs post transfection cells were stressed with arsenite (0.2mM) for 1hr before medium was replaced for the recovery period (2.5hrs). Cells were fixed with 4% paraformaldehyde for 10min before analysis for GFP fluorescence. For the fibroblasts derived from ALS patients, cells at a density of 3×10^5^ cells/ml were seeded on 12cm^2^ coverslips coated with collagen. Control or Nb9 expressing retrovirus was applied onto the medium and cells were centrifuged at 3500rpm for 90min at 37^°^C. 48hrs post-infection cells were treated with 0.5mM arsenite and allowed to recover in fresh medium for 1hr before immunostaining.

### Immunofluorescence staining

Cells were plated on coverslips in a 12-well plate and the day after were transfected as indicated. Cells were washed with PBS, fixed with 3.7% paraformaldehyde in PBS for 15min, then permeabilized with 1% Triton-X100 in PBS at RT for 10min. Cells were blocked with 1% goat serum in 0.05% PBS-Tween for 30min then incubated with primary antibody (diluted in 1% goat serum in 0.05% PBS-Tween) for 16hrs. After washes in 0.05% PBS-Tween, coverslips were incubated with FITC - or TRITC - conjugated donkey-anti-mouse or donkey-anti-rabbit secondary antibodies (diluted in 1% goat serum in 0.05% PBS-Tween) at RT for 1hr. DNA was stained with DAPI (1:20000) at RT for 1min. After mounting with ProLong Gold Antifade reagent (Molecular Probes), images were acquired with a Zeiss Axio Imager Z2 microscope with a 40X Plan Neofluar 1.3 NA (oil) or a 63X Plan Apochromat 1.4 NA oil objective and a scMOS ZYLA 5.5 camera controlled by the MetaMorph software (Universal Imaging, Roper Scientific). Images were treated with ImageJ or FIJI.

### Live-cell microscopy

For live cell microscopy, images were acquired using a 40X LUCPLFLN 0.6NA RC2 lens on an Inverted Olympus IX83 microscope controlled by Metamorph software and equipped with a full-enclosure environmental chamber heated to 37°C with 5% CO2 and an 1 ZYLA 4.2 MP scMOS camera. Frames were recorded every 3min over 1hr for SGs assembly and every 30min for 16hrs for SGs recovery. Parental and NEDP1 knockout U2OS cells were followed during the same time-lapse. Images were imported as a sequence and were analyzed using the Fiji software. Fiji was also used for brightness adjustment, cropping, creating scale bars and insets.

### Fluorescence recovery after photobleaching

Parental or NEDP1 knockout cells U2OS cells were grown on µ-Slide 8 Well – ibiTreat (ibidi GmbH, Gräfelfing, Germany). Immediately before imaging, cells were treated with 0.2mM arsenite and placed on a heated chamber at 37°C with 5% CO2. Imaging and photobleaching were performed with an apochromat ×100 oil objective using a Nikon TIRF PALM STORM inverted microscope. A 488-nm laser was used to photobleach G3BP1 inclusion after 1hr and 2hrs of arsenite treatment. Immediately after bleaching, the images were collected every 50msec for a total of 300 frames as a post-bleached sample. The fluorescence intensities of post-bleached G3BP1 inclusions (BL) were individually measured. In the meantime, the fluorescence intensities of nonbleached G3BP1 inclusions were measured as a reference control (REF). Also, the fluorescence intensities of background were measured as a background control (BG). To obtain corrected values (corr1), (BG) values were subtracted from Bleach (BL) and Reference (REF): BL_corr1(t) = BL(t) – BG(t); REF_corr1(t) = REF(t) – BG(t) then the corrected Bleach values were normalized to corrected Reference values: BL_corr2(t) = BL_corr1(t) / REF_corr1(t). The mean pre-bleach intensity was used to normalize the corrected bleach value: BL_corr3(t) = BL_corr2(t) / BL_corr2(pre-bleach). The mean pre-bleach value represents the maximum intensity to which the bleached region could possibly recover. The percentage of mobile fraction and the half time of recovery were obtained by using the curve fitting function/ exponential recovery in FIJI.

### Microtubule dynamics

For the time-lapse acquisition of EB1-mcherry signal, parental and NEDP1 knockout U2OS cells stably expressing EB1-mcherry were imaged by Nikon inverted microscope coupled to the Andor Dragonfly spinning disk. Images were acquired using an EMCCD iXon888 Life Andor cameras (pixel size: 0.11μm) with a 100x oil-immersion objective (Plan Apo lambda 1.45 NA 0.13mm WD). Images were acquired every 0.3sec during 45sec at 37°C in a thermo regulated atmosphere. Signal was then tracked in each individual cell using Matlab and the PlusTipTracker open source software package with the following parameters (Gaumes et al., 2016): *σ*1 = 1 / *σ*2 = 4 / K = 3 / Search radius range = 1–5 pixels / Minimum sub-track length = 3 frames / Maximum gap length = 30 frames / Maximum shrinkage factor = 1.5 / Maximum angle forward = 30°/ Maximum angle backward = 10°/ Fluctuation radius = 2 pixels. Results tables were then exported and further analyzed using Excel.

### FACS analysis

U2OS cells (6cm dishes) were transfected with either empty or GFP expressing pcDNA3 construct (1[g). 48hrs post-transfection cells were harvested, processed on a Aria IIU Becton Dickinson cytometer and data were analysed by FACSDiVa software.

### Cell viability-Clonogenic assays

For cell viability assays, cells were seeded in 96-well plates (500 cells/well) and treated as indicated. Cell viability was measured in triplicates using the CellTiter-GloR Luminescence assays from Promega or by counting live cells using Countess 3 (Invitrogen) according to manufacturer’s instructions. For the clonogenic assay, cells were seeded in 6-well plates (20000 cells/well) and treated as indicated. Plates were washed fixed with methanol for 20min at RT, dried, colonies were stained with Giemsa-stain and counted.

### Western blot analysis

Proteins were resolved on precast Novex (Invitrogen) SDS polyacrylamide gels and transferred onto nitrocellulose membrane (GE Healthcare). Membranes were washed once with PBS and blocked with 5% milk solution (PBS, 0.1% Tween-20%, 5% skimmed milk) for 1hr at room temperature with gentle agitation. Then membranes were washed 3 times with PBST (PBS with 0.1% Tween-20) for 10min each and incubated with primary antibodies (diluted in PBS, 0.1% Tween-20, 3% BSA, 0.1% NaN3) overnight at 4°C or for 3hrs at room temperature with gentle agitation. After washing 3 times with PBST for 15min, membranes were incubated with secondary antibodies diluted in 5% milk solution for 1hr at room temperature with gentle agitation. Then membranes were washed 3 times with PBST and 2 times with PBS for 15min each before soaked in ECL Western Blot Detection solution (Amersham) and exposure to Medical Films (Konica Minolta).

### Mass spectrometry

Di-Glycine motif peptide identification was performed by Cell Signaling Technology, following UbiScan protocols and instructions (Cell Signaling Technology) using Ubiquitin Branch Motif Antibody (K-ε-GG) [3925. Peptides were loaded onto 10cm x 75μm PicoFrit Capillary column packed with Magic C18 AQ reversed-phase resin. The column was developed with a 90min linear gradient of acetonitrile in 0.125% formic acid delivered at 280nL/min. MS parameters settings: MS run time 96min, MS1 Scan Range (300.0-1500.00), Top 20 MS/MS (Min Signal 500, isolation width 2.0, normalized coll. energy 35.0, activation-Q 0.250, activation time 20.0, lock mass 371.101237, charge state rejection enabled, charge state 1+ rejected, dynamic exclusion enabled, repeat count 1, repeat duration 35.0, exclusion list size 500, exclusion duration 40.0, exclusion mass width relative to mass, exclusion mass width 10 ppm). MS/MS spectra were evaluated using SEQUEST 3G and the SORCERER 2 platform from Sage-N Research (v4.0, Milpitas CA). Searches were performed against the most recent update of the NCBI human database with mass accuracy of +/-50ppm for precursor ions and 1Da for product ions. Results were filtered with mass accuracy of +/-5 ppm on precursor ions and presence of the intended motif (K-ε-GG). LTQ-Orbitrap Velos was used (Thermoscientific). A 5% default false positive rate was used to filter the SORCERER results.

### PARP-1 site-directed mutagenesis

All KR PARP-1 mutants were generated by site directed mutagenesis and sequences were verified by automated sequencing.

### Imaging of C. elegans

For imaging of stress granules, GFP-GTBP-1 expressing young adults (L4) were placed on a fresh NGM seeded plate and for heat shock, plates were incubated at 30oC for 4hrs.

For the arsenite treatment, unseeded NGM plates were soaked with sodium arsenite to achieve a final concentration of 10mM. Arsenite coated plates were then seeded with bacteria and young adults were placed on these plates for 2hrs. After treatment, animals were mounted on 2% (wt/vol) agarose pads and immobilized using 25mM Levamisole in M9 buffer. Images of the anterior region of the worm consisting of pharynx were captured using a fluorescence microscope (Zeiss AxioImager Z2) with a Plan-Apochromat 100x/1.40 Oil objective using ZEN software (version 3.4.91, Blue edition). Representative images are presented following max-project of Z-stacked images using maximum intensity projection type with Fiji Image J software (Version 2.3.0/1.53f). Stress granules were counted using cell counter plugin. The size of SGs was measured using ImageJ.

### Swimming behaviour (Motility) assay

The protocol described in (31) was employed. Young adults (L4) were treated overnight with 5mM paraquat (oxidative stress) in NGM seeded plates. Three worms were placed in a drop of 5μl of M9 buffer inside the ring drawn using a hydrophobic PAP pen on microscopic slide. Videos were recorded at 22 frames/sec for 30sec and analysed with the CeleST software in Matlab R2019b (version 9.7.0.1216025; Mathworks, Inc., Natick, MA, USA) (34). Worms were randomly selected for analysis and the number of body waves initiated from either the head or tail per minute are presented as wave initiation rate. Swimming rates for at least 15 worms per genotype from 3 repeats are presented.

### Statistical analysis

Leica LAS (FRAP analysis), SEQUEST 3G and the SORCERER 2 (mass spectrometry analysis), GraphPad Prism, PlusTipTracker, ImageJ and Cytoscape were used. For SG assembly/disassembly process, approx. 100 cells randomly selected per condition were scored for the presence of SGs. Statistical significance was calculated by Mann-Whitney test; ns: non-significant; *p < 0.05; **p < 0.01; ***p < 0.001. In all experiments n values represent the number of independent experiments, as indicated in figure legends.

