## Supplementary Figures for "Targeting the deNEDDylating enzyme NEDP1 to ameliorate ALS phenotypes through Stress Granules dissolution"

S1

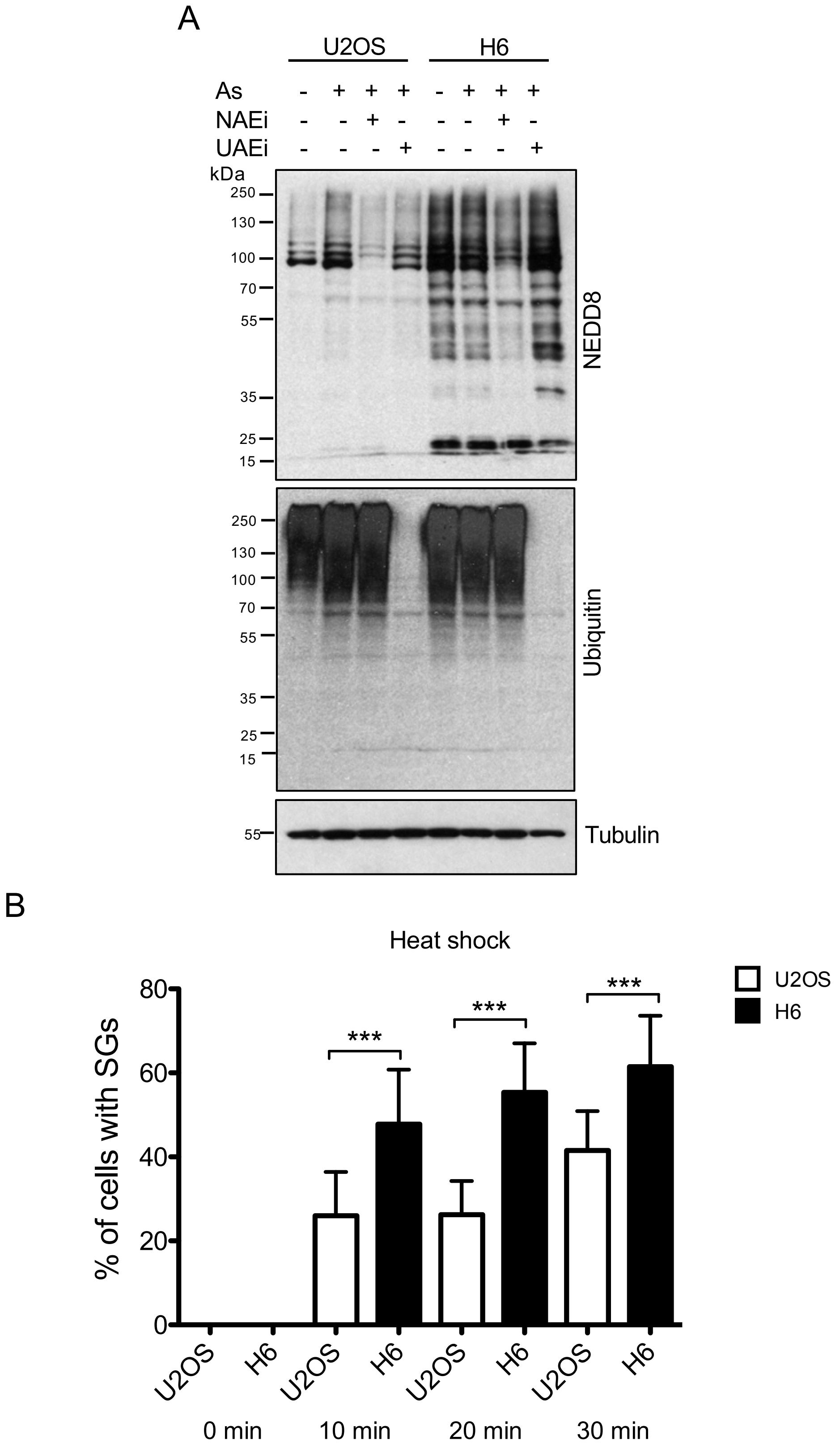


**Figure S1. The NEDD8 response to sodium arsenite depends on the canonical NEDD8 pathway. NEDP1 knockout accelerates SG formation in resposne to heat shock. Related to Figure 1.**

A. Parental and H6 NEDP1 knockout U2OS cells were pre-treated with NAEi and UAEi inhibitors (0.5μM, 4hrs) before cells were exposed to As (0.5mM, 1hr). Cell extracts were used for western blot analysis with the indicated antibodies. B. Parental and H6 cells stably expressing GFP-G3BP1 were exposed to heat shock (43°C) and SG formation was monitored by GFP fluorescence at the indicated time points. Quantitation was performed as in Fig 1B.

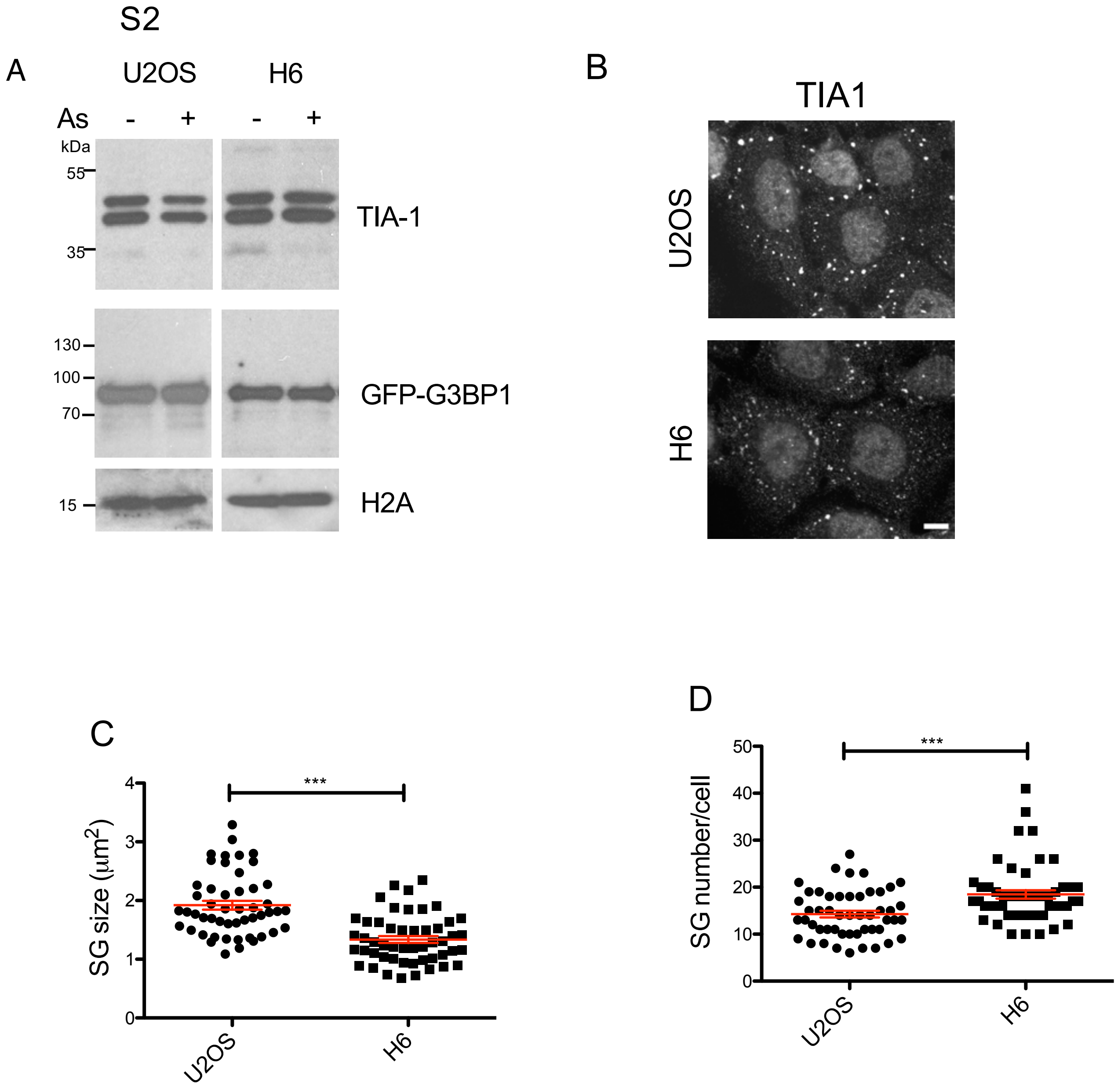


**Figure S2. NEDP1 knockout does not affect the levels of GFP-G3BP1 and decreases the size of SGs. Related to Figure 1.**

A. Parental and H6 NEDP1 knockout U2OS cells stably expressing GFP-G3BP1 were exposed to As (0.5mM, 1hr). Cell extracts were used for western blot analysis with the indicated antibodies. Parental and H6 NEDP1 knockout U2OS cells were treated with As (0.5mM, 1hr), before staining for TIA1 (B). The size (C) and number (D) of SGs were calculated as described in Star Methods.

S3

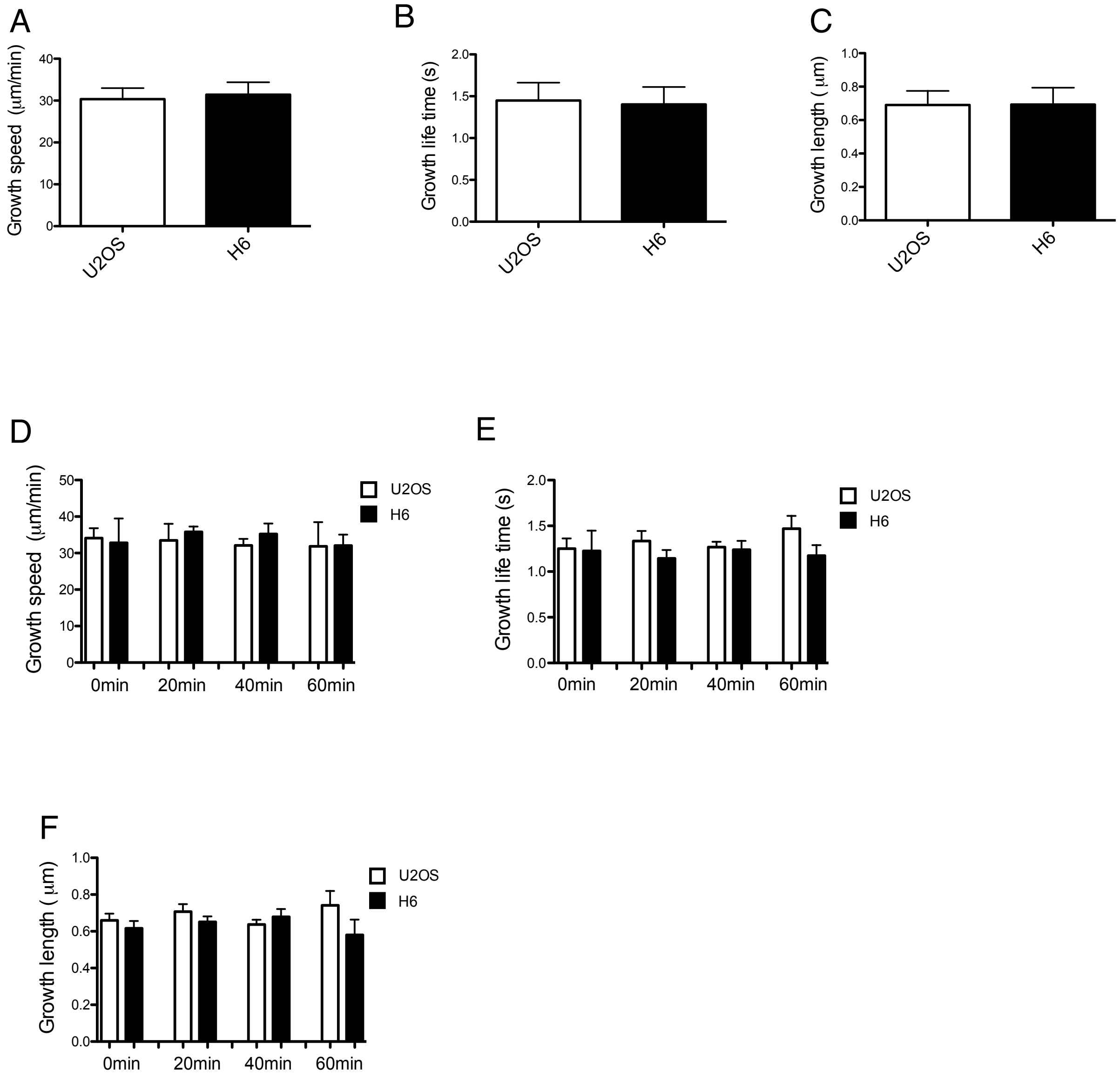


**Figure S3. NEDP1 knockout does not affect microtubule dynamics. Related to Figure1.**

Graphs represent tracks for (A) growth speed, (B) growth life time and (C) growth displacement (n=20 cells). Scale bar, 10µm. (D-F). Parental and H6 U2OS cells stably expressing EB1-mcherry were treated with 0.2mM As. Graphs represent tracks for (D) growth speed, (E) growth life time and (F) growth displacement for indicated time points. (n=3-4 cells).

S4

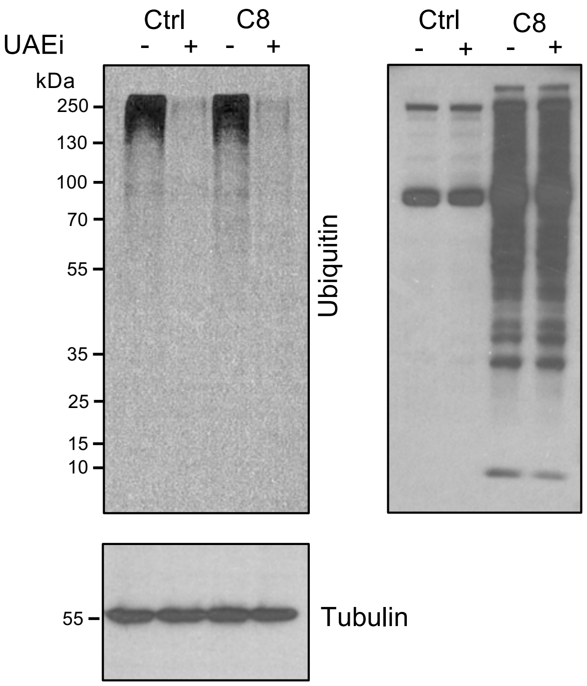


**Figure S4. Effect of UAEi on NEDD8, Ubiquitin pathways. Related to Figure 4.**
Parental and NEDP1 knockout HCT116 cells (C8) were either untreated or treated with UAEi (0.5µM, 5hrs) and cell extracts were analysed by western blotting with the indicated antibodies.

S5

A

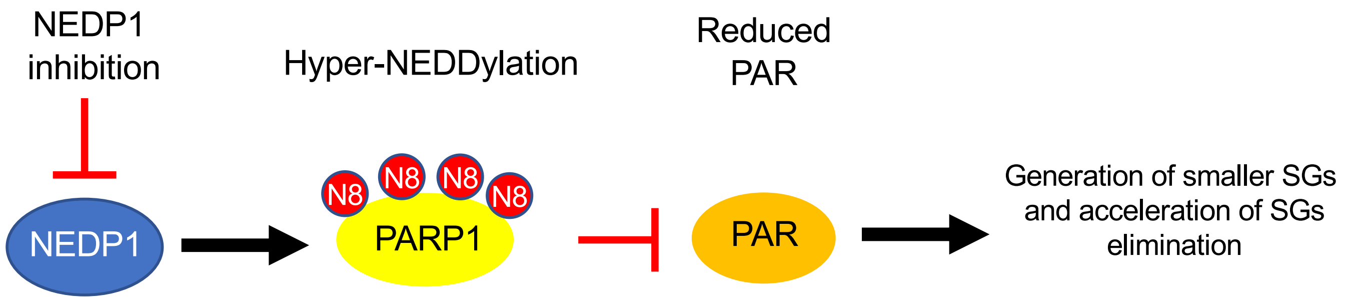


B

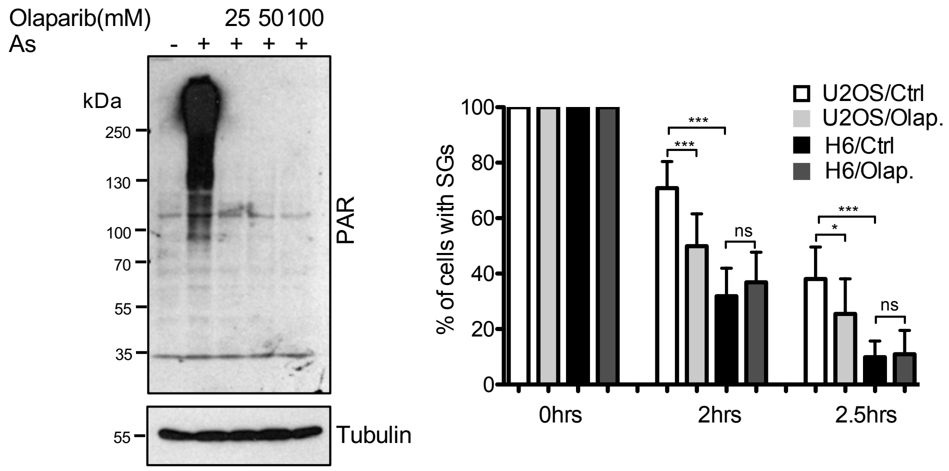


**Figure S5. NEDP1 and PARP-1 are functionally linked for the control of SGs elimination. Related to Figure 5.**
A. Model for the role of NEDP1 in PARP-1 activity control. Inhibition of NEDP1 causes the hyper-NEDDylation of PARP-1 that results in reduced levels of PAR production. This causes the generation of smaller SGs and accelerates their elimination during the recovery process. B. (Left panel). U2OS were pre-treated for 1hr with the PARP-1 inhibitor Olaparib followed by As treatment (0.5mM, 1hr). Cell extracts were used for western blot analysis with the indicated antibodies. (Right panel). Parental and H6 NEDP1 knockout U2OS cells stably expressing GFP-G3BP1 where pre-treated with Olaparib (25mM, 1hr) before As treatment as before, followed by recovery. SGs were monitored by GFP fluorescence. The graph represents the percentage of cells with SGs during the recovery period. For each condition the values where relative to the 0 time point (start of recovery). Values are the mean ± SD of 3 independent experiments.

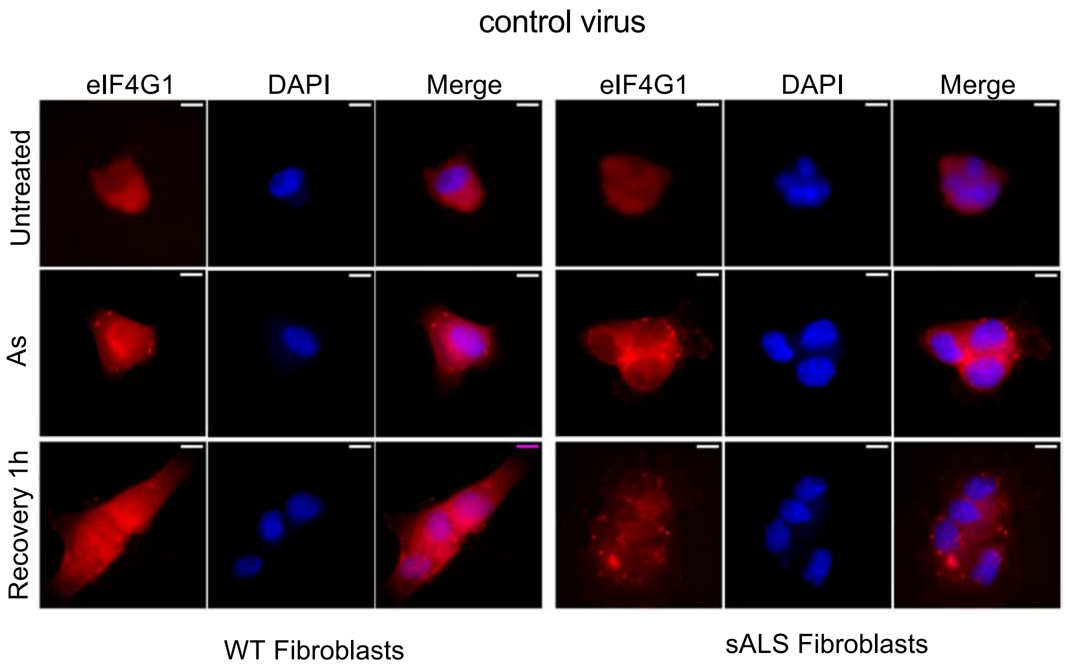


**Figure S6. Retrovirus infection does not affect SG dynamics. Related to Figure 6.** Experiment performed as in Figure 6F using instead a control (empty pMXs vector) retrovirus for infection.

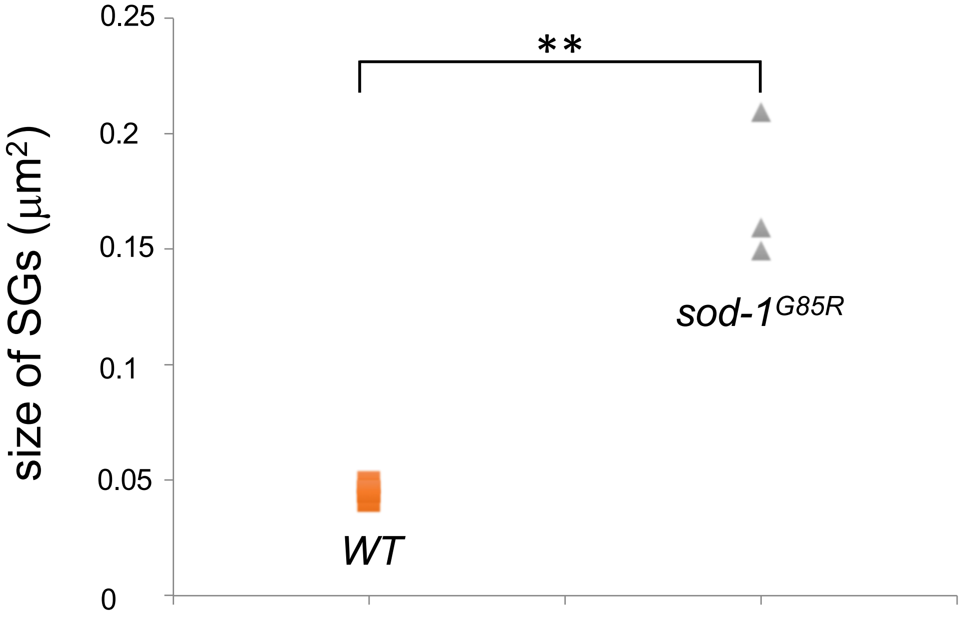


**Figure S7. Increased size of SGs in the *sod1*$^{G85R}$ mutant. Related to Figure 7.**
The size of SGs induced upon As treatment in the indicated *C. elegans* background was calculated as described in Star Methods. Each dot represents the average size of 50 randomly selected SGs in one animal. Experiment was performed 3 times.
